# Adhesion-driven invasion: disentangling the interplay between cell-cell and cell-matrix interactions in cancer cell migration

**DOI:** 10.1101/2025.06.23.660988

**Authors:** Quirine J. S. Braat, Klara Beslmüller, Cornelis Storm, Erik H. J. Danen, Liesbeth M. C. Janssen

## Abstract

Metastasis proceeds through the dissemination of cells from the primary tumor into the surrounding extracellular matrix (ECM), initiating further spread throughout the body. The invasion of cancer cells into the ECM is significantly influenced by cell-cell adhesion, cell-matrix adhesion, and the generation of traction forces. However, the complex interplay between these different aspects makes it difficult to disentangle their roles in the invasive process, hampering our mechanistic understanding of collective cell invasion. Here, we combine integrin knockout experiments in Hs578T and 4T1 breast cancer cell lines with a computational cellular Potts model to elucidate the biophysical mechanisms underlying invasive cell behavior. By tuning the cell-cell and cell-matrix adhesion parameters in our computational model, we establish a quantitative mapping with the experiments. Our work reveals that strong cell-matrix interactions promote invasion, while strong cell-cell adhesion promotes the formation and dissemination of multicellular clusters. Moreover, our model delineates a threshold for invasion and predicts that tumor morphology – particularly the number of branches from the primary tumor – correlates with its invasive potential, suggesting that morphological tumor features may serve as a proxy for metastasis. Our approach highlights the importance of combining biological experiments with computational and predictive modeling, offering new insights into the mechanisms driving cancer cell migration.

## 1. INTRODUCTION

As cancer cells disseminate from the primary tumor, their ability to invade new tissues relies heavily on directed migration through the extracellular matrix (ECM) [1, 2]. The ECM not only provides structural support but also serves as a dynamic environment that guides or impedes cell movement. Effective migration depends on several biophysical factors, including cell-ECM adhesion, traction force generation, ECM remodeling, and cell-cell interactions [3, 4]. In vivo, tumor cells often migrate along aligned collagen fibers or preexisting tracks within the ECM [5], with collagen bundles frequently oriented perpendicular to the tumor boundary [6, 7]. However, excessive ECM density can hinder invasion [8], highlighting the importance of finely tuned adhesion and matrix composition in supporting invasive migration. Adhesion to ECM components such as collagen is therefore a key driver of invasive migration.

At the single-cell level, migration is driven by actin-based protrusions at the leading edge, followed by cell-matrix adhesion and actomyosin-mediated contraction of the cell body [9]. These processes are coordinated through integrins – transmembrane receptors that physically link the intracellular actin cytoskeleton to the extracellular matrix [10, 11]. Integrins are obligate *αβ* heterodimers, with both subunits required for effective force transmission and adhesion. In collagen-rich ECM, migration depends on integrin-mediated adhesion and the generation of traction forces [12]. Disruption of integrin function, such as through inhibition of *α*v*β*5 [13] or *β*1 [14], can suppress invasion. It should be noted, however, that the outcome is cell type–dependent; for instance, *β*1 depletion may enhance migration via a switch in migration mode [15]. Importantly, in many cell types, *α*2*β*1 is abundantly expressed and plays a central role in collagen binding and invasive behavior [10].

In addition to cell-ECM adhesion, migration is also regulated by adhesion between neighboring cells. Cadherin-based junctions enable cells to form stable intercellular contacts and maintain tissue cohesion [16]. This mechanical coupling is particularly important during collective migration, where cells move as coordinated groups [17–19]. In epithelial and cancerous tissues, E-cadherin is a key mediator of stable cell-cell adhesion, promoting the formation of cohesive cell clusters [20]. Loss of E-cadherin, often accompanied by a switch to N-cadherin expression, is associated with a more fibroblastic, motile, and invasive phenotype [21]. Emerging evidence suggests crosstalk between cadherin- and integrin-based adhesions, indicating a coordinated regulation of cell-cell and cell-matrix interactions during invasion [14, 22], but a complete mechanistic understanding is still lacking.

To better understand and predict the roles of cadherins and integrins in cancer cell invasion, computational modeling has become an essential tool [23, 24]. A variety of models have been developed to study how cell-cell and cell-matrix interactions influence migration of cells into the ECM [25–32]. While the interplay between cell-cell adhesion and ECM confinement has been explored [25, 33–35], less is known about how cell-matrix adhesion shapes cellular responses to the ECM. For instance, Kang *et al*. [25] demonstrated that cell motility and ECM density are key determinants of invasiveness, but their computational model does not explicitly incorporate adhesive interactions between cells and the ECM. Conversely, models that include detailed cell-matrix dynamics often focus on single-cell behavior [36–38], limiting their ability to capture the complex interplay between cell-cell and cell-matrix adhesion observed in multicellular spheroid experiments. Moreover, few models are designed for direct, quantitative comparison with experimental data across different cell lines, which restricts their capacity to generate mechanistic insights that are experimentally testable.

Here, we address this gap by developing a computational model in combination with integrin knockout experiments in two breast cancer cell lines, aimed at disentangling the interplay between cell-cell and cell-matrix adhesion during cell invasion. Our integrated approach enables direct comparison between simulated and experimentally observed invasion behaviors, allowing us to dissect how specific adhesion mechanisms influence collective invasion dynamics. By systematically tuning cell-cell and cell-matrix adhesion parameters to reflect different experimental conditions, we establish a quantitative mapping between model predictions and experimentally observed invasion behaviors. Furthermore, our model predicts that tumor morphology – particularly the number of invasive branches – correlates with overall invasiveness, suggesting that morphological features may serve as proxies for metastatic potential. Finally, our model highlights the critical role of traction forces in initiating invasion and suggests that ECM binding modulates the traction forces that cells generate. Together, these findings demonstrate the value of combining experimental and computational methods to uncover the biophysical principles of cancer cell invasion.

## 2. METHODS

### A. Experimental methods

4T1 mouse breast cancer cells and Hs578T human breast cancer cells were used, that differ in their cell-cell adhesion strength, with 4T1 displaying stronger cell-cell adhesion than Hs578T. These cell lines were genetically modified to express different levels of cellmatrix binding. For this, Hs578T cells were transduced with a third-generation lentiviral packaging system, expressing a conditional Cas9 vector and sgRNAs targeting ITGA2 and ITGB1. For 4T1, a vector was designed combining stable Cas9 and sgRNAs. Knock-outs were verified using Western blot and proliferation and adhesion to collagen was assessed. Wildtype and knockout cells were used in a 3D tumoroid model, in which tumoroids are printed in a pre-formed collagen gel. Their invasion and migration into the surrounding ECM was monitored with confocal microscopy. Full details of all experimental methods can be found in the SI.

### B. Computational Model

To investigate the interplay between cell-cell and cell-matrix interactions in cell migration, we developed a computational model based on the cellular Potts (CP) formalism [39, 40]. Briefly, our model consists of a static, fibrous ECM structure and a dynamic primary tumor core (see Figure 1A). We opted for a quasi-two-dimensional setup for computational simplicity, as it also allows us to study the competition between cell-cell adhesion and different cellmatrix interactions during migration in a tractable manner. Our simulations thus represent a plane section of the tumoroid. The model is implemented in CompuCell3D [41].

**Figure 1.**
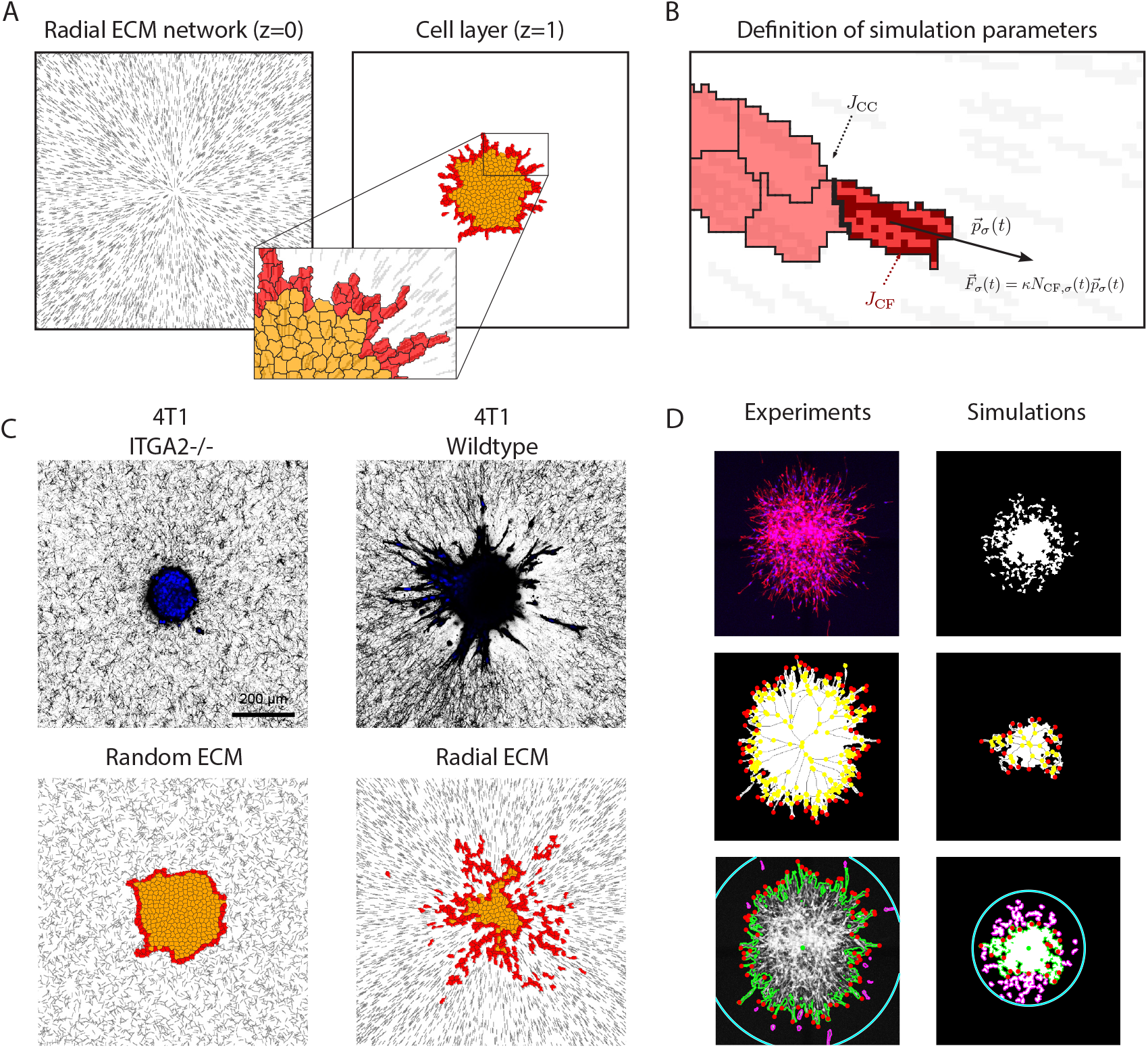
Computational Cellular Potts Model and tumoroid analysis by in-house Matlab script. **A**. Representation of the primary tumor and ECM fibers in the CP model. The fibers are represented as lines in the bottom layer (*z* = 0) and the primary tumor forms the top layer (*z* = 1). The tumor contains two types of cells, namely invasive cells (red) and noninvasive cells (orange). Invasive cells, in contact with ECM fibers, migrate into available space. **B**. Schematic representation of the simulation parameters. *J*_CC_ sets the cell-cell adhesion between cells (bold), *J*_CF_ sets the cell-matrix adhesion between cell and fiber pixels (darker tone). The force per ECM binding site is defined as *κ*. More details regarding the force of the cells is given in the main text. **C**. Tumoroids in random collagen network (left) and radial collagen network (right) for both experimental data (top) and computational data (bottom). **D**. Analysis of the experimental and simulation images with in-house Matlab script. From top to bottom, the input image, the black-and-white representation of the primary tumor after analysis, and the final analysis including the detached cells are shown.

Within our setup, the ECM fibers are represented as discrete fibers on a square lattice, similar to previous modeling work [30, 35, 42–45]. The orientation of these fibers is based on experimentally observed ECM networks (see Figure 1C). We consider two limiting cases, namely a random ECM and a radial ECM topology. We focus primarily on results obtained with the radial ECM topology; The random ECM topology is used as a benchmark in absence of any ECM remodeling.

The tumoroid is placed on top of the ECM fiber configuration (see Figure 1A). The primary tumor core contains two distinct cell types: nonmotile tumor cells and motile invasive cells. Both cell types possess the same specific surface and volume properties, reflecting the cortical tension and compressibility of the cell [46], respectively. In addition to these intracellular cell properties, the cells can adhere to other cells and to ECM fibers. We denote the cell-cell adhesion strength with *J*_CC_ and the cell-matrix adhesion strength with *J*_CF_. In the absence of either cell-cell adhesion or cell-matrix adhesion, we set the adhesion strengths equal to zero. When cells prefer to adhere to the ECM fibers, *J*_CF_ increases to promote cell-matrix binding. An increase in *J*_CC_, on the other hand, promotes cell-cell contacts such that cells are more strongly bound to each other. Thus, by adjusting the values of *J*_CC_ and *J*_CF_, we can modify the adhesion interactions with the extracellular matrix and other cells.

The motile invasive cells also interact with the ECM through force generation, allowing them to actively migrate outwards. Here we define invasive cells as those that form the boundary of the primary tumor and that are in contact with at least one ECM fiber (see zoom-in in Figure 1A). The traction force generated by the invasive cells is modeled with a self-alignment mechanism [47], similar to previous modeling works [48–51]. This alignment rule ensures that the invasive cells migrate radially outwards into the ECM. Moreover, we assume that the traction force scales with the number of ECM binding sites, inspired by [52], and a traction force per ECM binding site (*κ*). An overview of the cell-cell and cell-matrix interactions that are included in the computational model are schematically shown in Figure 1B.

We studied the migratory cell behavior as a function of the traction force per ECM binding site (*κ*), the cell-cell adhesion strength (*J*_CC_), and the cellmatrix adhesion strength (*J*_CF_). An overview of the complete set of simulation parameters, including the definition of the Hamiltonian in the CP formalism, is provided in the Supplementary Material.

### C. Analysis

To quantify the invasiveness and morphologies of our tumor spheroids, we employ consistent analysis methods for both our tumoroid experiments and the CP simulations. To this end, we use an in-house Matlab script adapted from Hou *et al*. [53].

For experimental data, scanning confocal z-stacks of the actin cytoskeleton were projected using the standard deviation in the z-direction. A Gaussian filter with a narrow kernel was used to remove small fluctuations from the projected image. An adaptive threshold was used to separate the tumoroid from the background. Figure 1D (left) shows snapshots of the procedure for the experiments. The bottom panel shows the different measurable invasion parameters. The primary tumor core is outlined with the green line and allows for extraction of the tumoroid area *A* and perimeter *P*. The invasive radius (cyan line) is defined as the distance traveled by the cell farthest from the centroid. When the tumor is non-invasive, that is, no cells detach from the tumor core, the invasive radius is set to 0. Skeletonization of the foreground mask, with the endpoints of the resulting skeleton serving as the branch points, allows us to detect the number of branches of migrating cells (red dots in Figure 1D).

For the simulations, the same Matlab analysis script was used (see Figure 1D (right)). Since the simulation snapshots are black-and-white images, the adaptive threshold method was not applied. The invasive radius extracted from our CP simulation data can be used as a direct comparison with the experiments. To provide additional information on the size distribution of the detached clusters of cells, which is more difficult to extract from the experiments, we also detect the number of detachments in simulation (outlined with magenta).

The morphology of the primary tumor is quantified by the complexity, defined in terms of the tumoroid perimeter and area as [53]

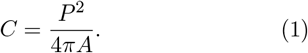

This dimensionless number characterizes the roughness of the tumor shape. Rough tumors have a relatively large surface area and are therefore characterized by a large complexity, whereas spherical tumors have a complexity *C* = 1.

We additionally extract information about the shape of individual cells. This analysis can only be performed for our simulation data, however, as the experimental resolution is insufficient to reliably measure individual cell morphologies. We quantify single-cell shapes in terms of the shape index [54], defined as 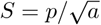 where *p* and *a* are the perimeter and area of a given cell, respectively. To correct for CP lattice artifacts in the determination of *p*, we follow the analysis procedure of Magno *et al*. [55].

## 3. RESULTS

### A. Cell-cell adhesion promotes collective invasion

To characterize the cell-cell adhesion strengths of the two different cell lines (Hs578T and 4T1 cells), we injected both cell lines as 3D tumoroids into preformed Collagen I matrices and measured their invasive properties. We studied the cell detachment from the tumoroid and migration into the surrounding ECM using a maximum projection of 5 z-slices of confocal microscopy images, as shown in Figure 2.

**Figure 2.**
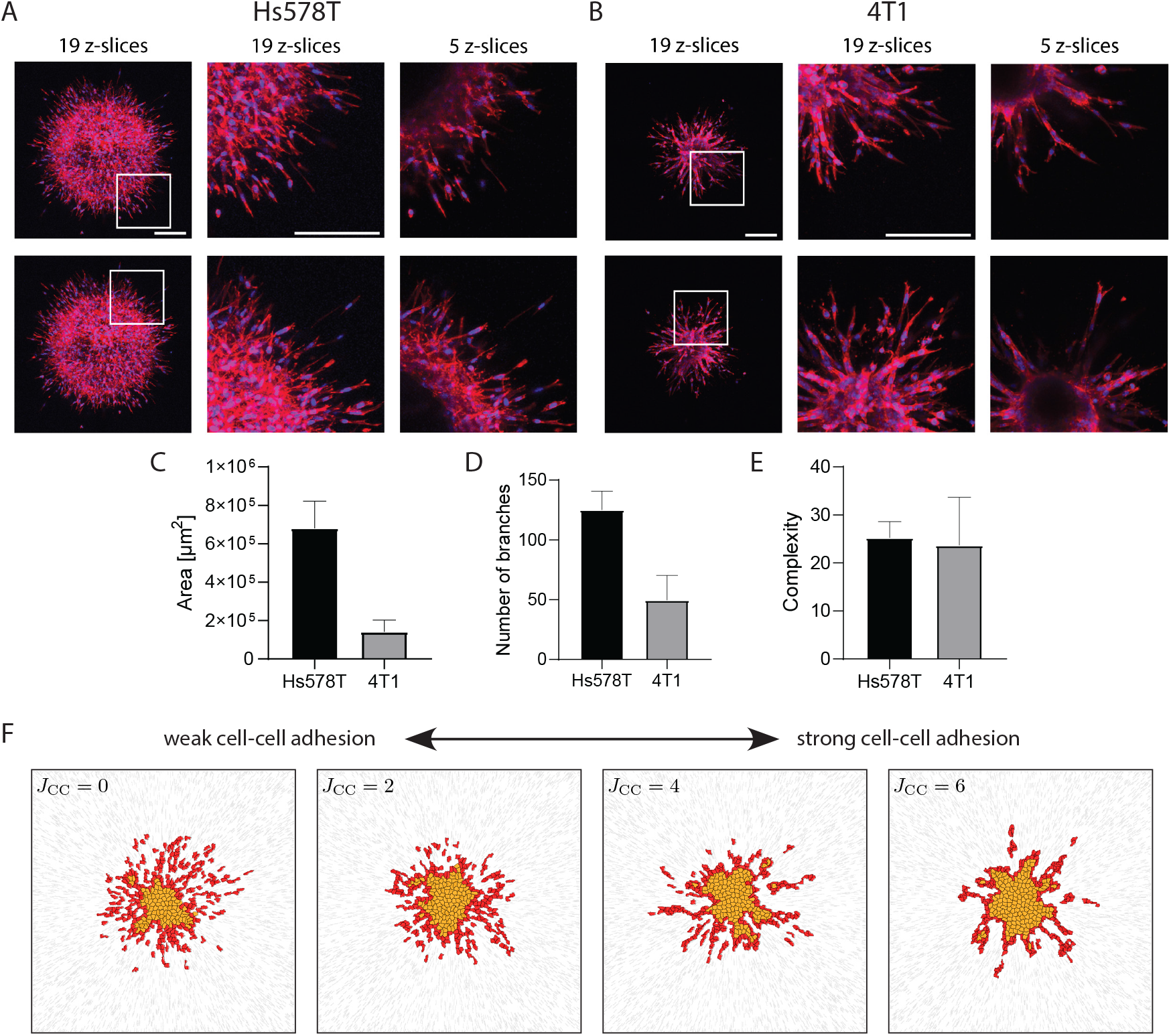
Migration modes of cells detaching from a tumoroid depend on the cell-cell adhesion strength. **A.** Maximum projections of representative confocal images of Hs578T tumoroids detaching from tumoroid as individual cells. Tumoroids were stained with Phalloidin (actin, red) and Hoechst (nuclei, blue). Scalebar: 400 µm. **B**. Maximum projections of representative confocal images of 4T1 tumoroids invading the ECM collectively. Tumoroids were stained with Phalloidin (actin, red) and Hoechst (nuclei, blue). Scalebar: 400 µm. **C**. Quantifications of tumoroid area for wildtype Hs578T cells and wildtype 4T1 cells. Mean *±* SD of three experiments with each a minimum of 4 tumoroids each are shown. **D**. Number of branches from tumoroid core for wildtype Hs578T cells and wildtype 4T1 cells. Mean *±* SD of three experiments with each a minimum of 4 tumoroids each are shown. **E**. Complexity of tumoroid for wildtype Hs578T cells and wildtype 4T1 cells. Mean *±* SD of three experiments with each a minimum of 4 tumoroids each are shown. **F**. Simulation snapshots with increasing cell-cell adhesion strength from no cell-cell adhesion (left) to strong cell-cell adhesion (right).

Hs578T cells disseminated from the tumoroid in a dispersed manner, with individual detached cells at the migration front (Figure 2A). Invading Hs578T cells exhibited an elongated mesenchymal phenotype and used filopodial spike-mediated strategies for migration. Invading Hs578T cells were not found in close contact with neighboring cells, indicating low cell-cell adhesion.

In contrast, detaching 4T1 cells moved in large groups and used collective invasion to migrate through the ECM (Figure 2B). These large clusters consisted of multicellular strands with elongated cells in direct contact with each other, suggesting stable cell-cell junctions. Overall, this more collective nature of 4T1 cells suggests relatively strong cell-cell adhesion.

To corroborate the differences in cell-cell adhesion strengths, we quantified the E- and N-cadherin expression in both cell lines. Figure S2 shows that Hs578T cells do not express E-cadherin but do express N-cadherin, whereas 4T1 cells show high Ecadherin expression and no N-cadherin expression. These findings, together with the above observations, confirm that cell-cell adhesion is effectively absent in Hs578T cells, and strongly present in 4T1 cells.

To further characterize the invasiveness of the tumoroids, we measured the tumoroid area, complexity, and number of branches of three independent experiments. The complexity of the tumoroid reflects the geometric intricacy of the outer surface of the tumor, with a higher value indicating more irregular shapes and a lower value indicating smoother surfaces. Our results show that Hs578T tumoroids have a larger area compared to the 4T1 tumoroids (Figure 2C), indicating that a lack of cell-cell adhesion promotes cell invasion. Similarly, Hs578T cells showed more branches inherent to the more invasive nature of the cells (Figure 2D). The tumoroid complexity remains similar for the two cell lines (Figure 2E), however, which stems either from a large tumoroid with many short strands (Hs578T) or a small tumoroid with long strands (4T1).

Differences in cell-cell adhesion were incorporated into the computational model using the cell-cell adhesion parameter *J*_CC_. Figure 2F shows four different simulation snapshots with increasing cell-cell adhesion strength from left to right. In the absence of cell-cell adhesion (*J*_CC_ = 0), cells migrate primarily as single cells. In contrast, strong cell-cell adhesion (*J*_CC_ = 6) results in the formation of collective strands. These observations, where cells with low cell-cell adhesion strength migrate as single cells and often invade further than cells with high cell-cell adhesion, are consistent with previous experiments as well [18]. Therefore, in the following sections, the migration patterns of Hs578T cells are compared with the simulations where *J*_CC_ = 0, while the migration patterns of 4T1 cells are compared with the simulations where *J*_CC_ = 4.

### B. Invasion of 4T1 and Hs578T cells into ECM is dependent on cell-matrix binding strength

Having characterized the differences in cell-cell adhesion strengths among the two cell lines, we next aimed to study the effect of cell-matrix interactions on the migration. To this end, we perturbed cell-ECM adhesion by knocking out ITGA2 or ITGB1 integrin subunits expected to prevent integrin *α*2*β*1-mediated interactions with the collagen network.

#### 4T1 cells

Stable 4T1 knockout cells were verified using Western blot (see Figure 3A). Knockout cells were further analyzed by testing the effect of the knockout on proliferation and collagen binding. 4T1 Itg*β*1^-/-^cells were not used since depletion of ITGB1 was shown to affect EMT status in these cells [15] and in our current experiments it reduced proliferation (Figure S3). Proliferation of 4T1 cells was not affected by depletion of ITGA2 (Figure 3B). mRNA analysis of all collagen receptors showed that *α*2*β*1 is the only collagen-binding integrin expressed in 4T1 (Figure S4. This was in line with the fact that deletion of ITGA2 attenuated 4T1 cell adhesion to collagen by 90.4% (Figure 3B) and Itg*α*2^-/-^cells were used in the following experiments.

**Figure 3.**
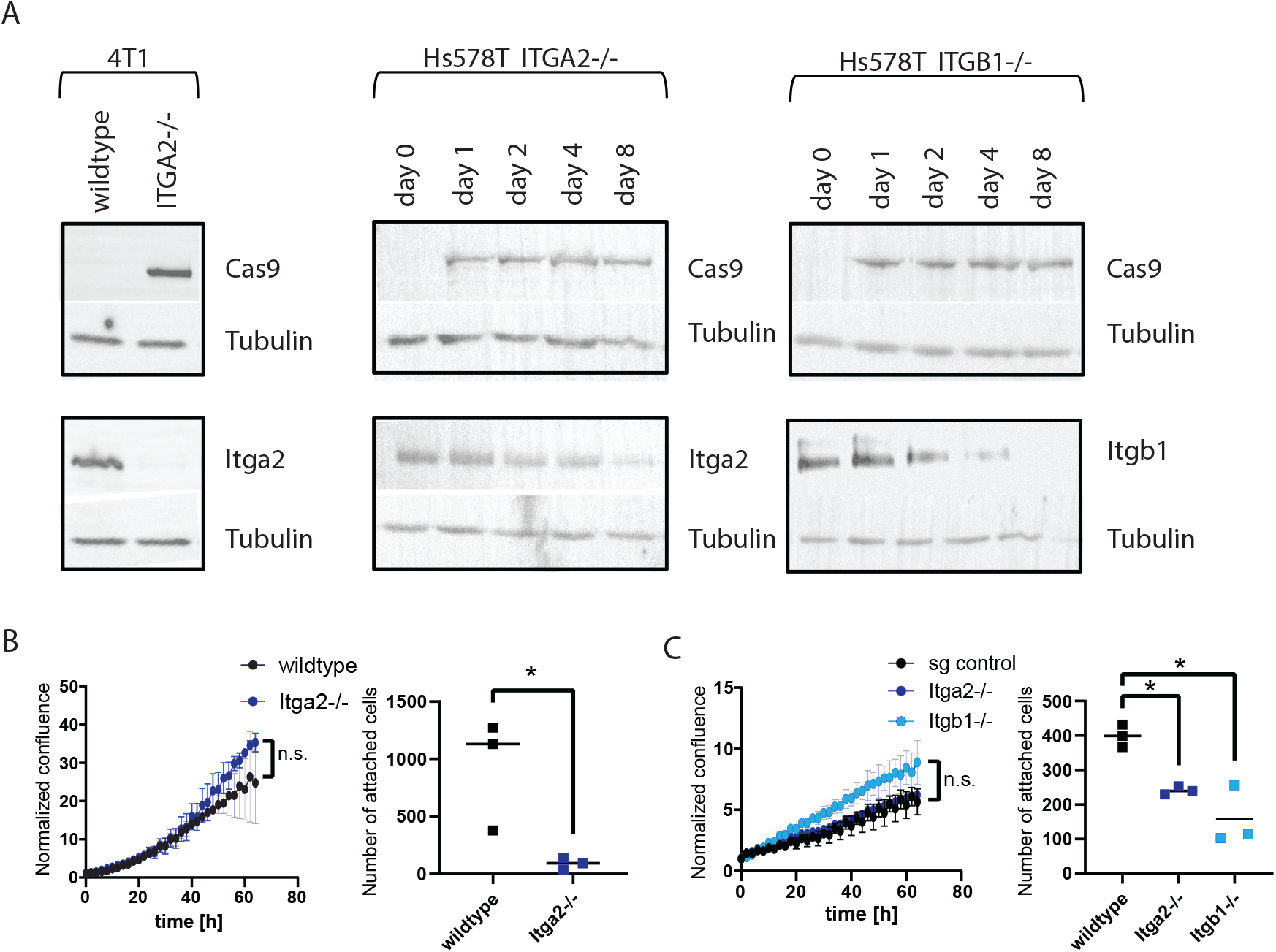
Characterization of integrin knockouts. **A**. Western blot showing stable knockout of ITGA2 in 4T1 and Doxycycline inducible knockout of ITGA2 and ITGB1 in Hs578T. Day indications refer to the amount of days that cells were treated with 1 µg/mL Doxycycline. **B**. Proliferation and adhesion to collagen of 4T1 knockout cells. Mean *±* SD of three experiments are shown with 12 technical replicates (for proliferation graphs) and a minimum of 4 technical replicates (for adhesion graphs). **C**. Proliferation and adhesion to collagen of Hs578T knockout cells. Mean*±* SD of three experiments are shown with 12 technical replicates (for proliferation graphs) and a minimum of 4 technical replicates (for adhesion graphs).

Cells were printed as 3D tumoroids in a collagen gel. For 4T1, Itg*α*2^-/-^tumoroids barely invaded the ECM (see Figure 4A) and hardly any branches were formed. The wildtype cells, on the other hand, formed many branches as cells showed collective strands that invaded far into the ECM. The reduction in invasion was quantified with the tumoroid area. Figure 4B shows that the quantified tumoroid area with Itg*α*2^-/-^cells was reduced by 60% compared to the wildtype tumoroids. In addition to the area, both the invasive radius and the complexity of the tumoroid significantly decreased with Itg*α*2^-/-^cells (see Figure 4C and S5, respectively).

**Figure 4.**
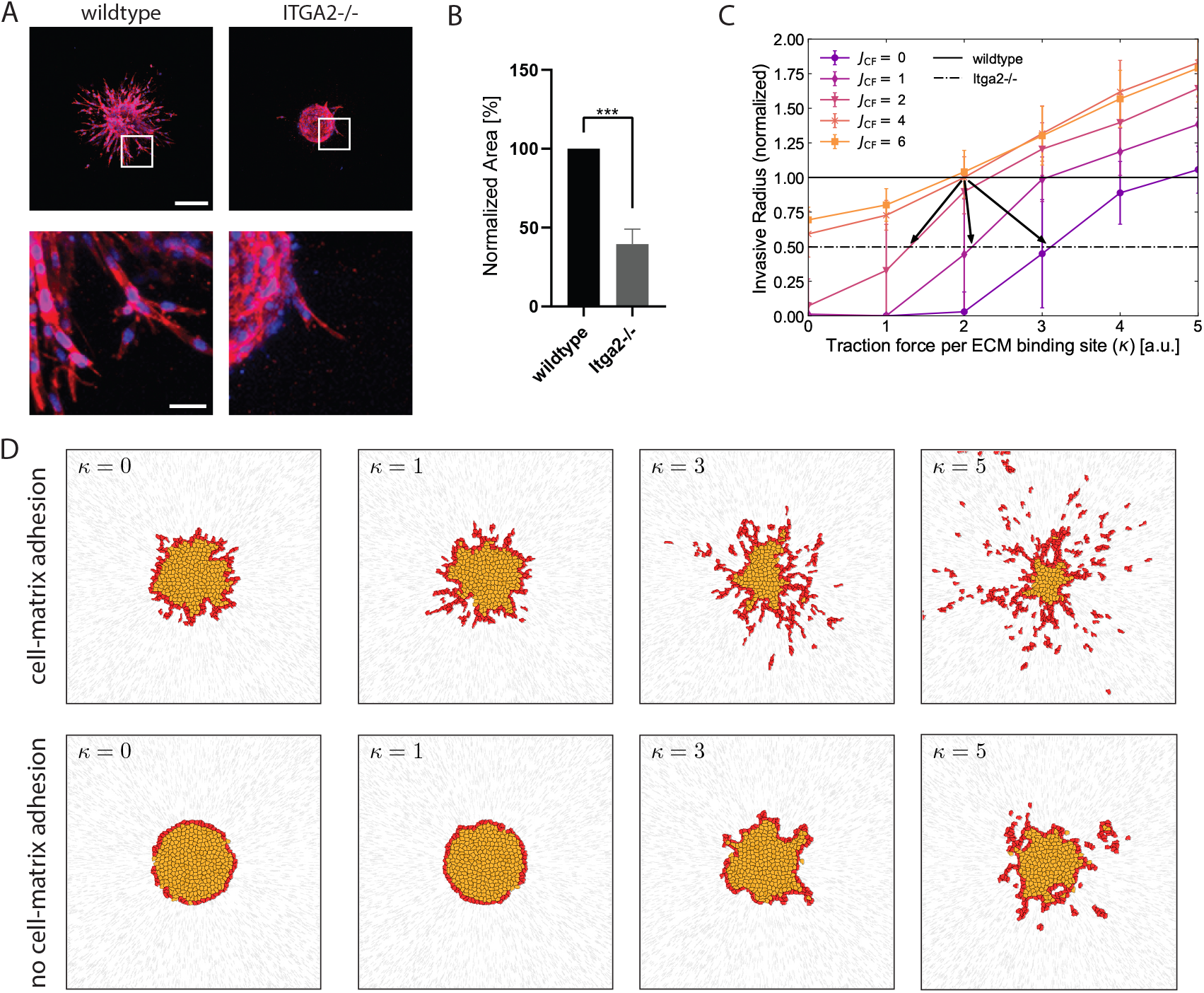
Invasion of cells with high cell-cell adhesion is dependent on cell-matrix binding in experimental data and modeling. **A**. Maximum projections of representative confocal images of 4T1 tumoroids of one representative tumoroid. Tumoroids were stained with Phalloidin (actin, red) and Hoechst (nuclei, blue). Scalebar: 200 µm. **B**. Area quantifications of 4T1 tumoroids showing mean *±* SD of three experiments with each a minimum of 4 tumoroids. Each experiment was normalized to area of wildtype=100%. **C**. Invasive radius as a function of the traction force per ECM binding site (*κ*) for various cell-matrix adhesion strengths *J*_CF_. The simulation data is normalized with respect to *κ* = 2 and *J*_CF_ = 4. The experimental data (horizontal lines) for the 4T1 cells are normalized with respect to the wildtype. **D**. Migration of cells with (*J*_CF_ = 4) and without (*J*_CF_ = 0) cell-matrix adhesion for the simulations with high cell-cell adhesion (*J*_CC_ = 4). The different snapshots show how the cells migrate as the traction force generated per ECM binding site (*κ*) increases.

In the simulations, we can mimic the change in cell-matrix interactions using two different parameters: the cell-matrix adhesion strength *J*_CF_, and the traction force parameter *κ*. Figure 4C and D show that increasing either the cell-matrix adhesion strength or the traction force generated by the cells promotes migration into the ECM. To identify where wildtype and Itg*α*2^-/-^cells lie in the parameter regime of the simulations, we compare our simulations for varying *J*_CF_ and *κ* with the experimental results. The intersection of the experimental data with the simulations in Figure 4C informs us about the relationship between the change in collagen binding for Itg*α*2^-/-^cells and the corresponding change in the cellmatrix interaction in terms of the simulation parameters.

The simulation proposes three potential scenarios that are indicated by the arrows. When Itg*α*2 is removed, cells can either 1) reduce the force they generate, while keeping their cell-matrix binding strength approximately the same (left arrow), generate the same force but significantly reduce the cell-matrix binding strength (middle arrow), or completely lose cell-matrix binding and generate more force through a potentially different migration mechanism (right arrow). The significant reduction in cell-matrix binding (Figure 3B, right panel) suggests that scenario 2 is most probable. To confirm this hypothesis, we use the second cell line (Hs578T), which exhibits a high, intermediate, and low cellmatrix interaction regime.

#### Hs578T cells

To study an intermediate collagen binding regime, we produced Doxycycline-inducible integrin knockouts in Hs578T cells. The Western blot results (Figure 3A) showed that Itg*β*1^-/-^was completely removed for Hs578T cells after Doxycycline treatment for 8 days. Itg*α*2^-/-^cells still had slightly remaining protein, even after 2 weeks of Doxycycline induction (data not shown). For all subsequent experiments with Hs578T cells, knockout was induced by treatment with Doxycycline for 8 days. Both Hs578T Itg*α*2^-/-^and Itg*β*1^-/-^cells led to impaired collagen binding (reduction of 39.7% for Itg*α*2^-/-^and 60.5% for Itg*β*1^-/-^), while the proliferation of these cells was not affected by the knockout (Figure 3C). Both knockout cell lines were printed as tumoroids in a collagen gel, similar to the 4T1 cells. The invasion of integrin-depleted cells from the tumoroid into the ECM was significantly reduced compared to the invasion of sg control cells (Figure 5A). Itg*α*2^-/-^tumoroids remained smaller in area compared to the sg control cells, while the Itg*β*1^-/-^tumoroids hardly invaded at all into the ECM. Quantification of the tumoroid area revealed that Itg*α*2^-/-^reduced the invasion by 44% and Itg*β*1^-/-^reduced the invasion by 85% (Figure 5B). In addition to the area, the number of branches formed from the tumoroid core was also significantly reduced by 23% for Itg*α*2^-/-^cells and by 76% for Itg*β*1^-/-^cells as compared to sg control cells (Figure 5C). The complexity of the tumoroid increased in Itg*α*2^-/-^cells compared to the sg control cells (not significantly, see Figure S5). However, with Itg*β*1^-/-^cells the complexity was significantly reduced (9 branches compared to 25, averaged over 3 experiments), as almost no branches formed and the tumoroid remained spherical.

**Figure 5.**
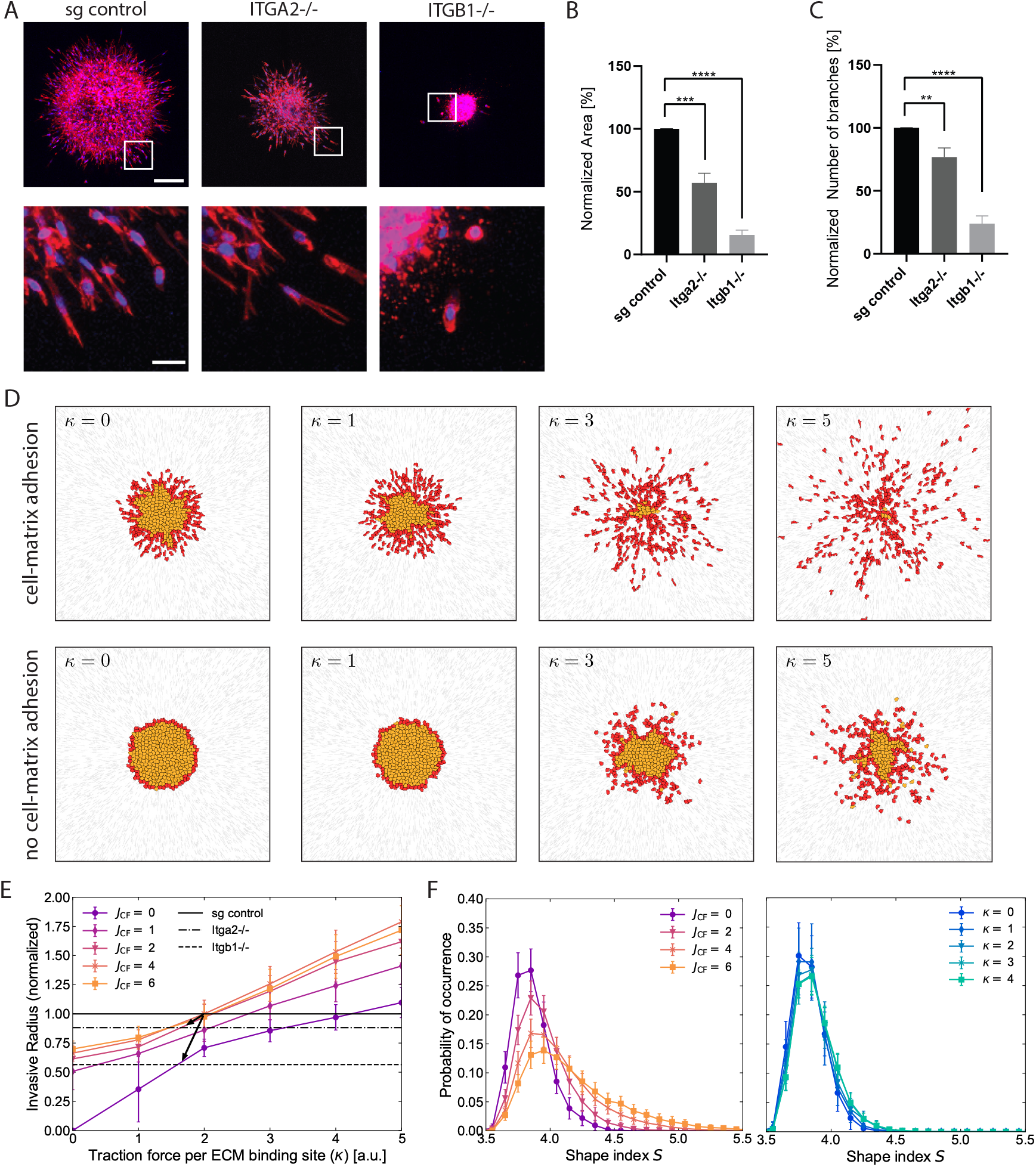
Invasion of cells with low cell-cell adhesion is dependent on cell-matrix binding in experimental data and modeling. **A**. Maximum projections of representative confocal images of Hs578T tumoroids. Scalebar: 200 µm. **B**. Area quantifications of Hs578T tumoroids showing mean *±* SD of three experiments with each a minimum of 4 tumoroids. Each experiment was normalized to area of sg control=100%. **C**. Number of branches forming from the core of Hs578T tumoroids showing mean *±* SD of three experiments with each a minimum of 4 tumoroids. Each experiment was normalized to area of sg control=100%. **D**. Migration of cells with (*J*_CF_ = 4) and without (*J*_CF_ = 0) cell-matrix adhesion for the simulations without cell-cell adhesion (*J*_CC_ = 0). **E**. Invasive radius as a function of *κ* for various cell-matrix adhesion strengths *J*_CF_. Results are normalized to *κ* = 2 and *J*_CF_ = 4 (simulations) or sg control (experimental). The conclusions remain similar for different normalization in the simulations (see Figure S7). **F**. Shape distribution of the invasive (red) cells. Shape distribution for fixed *κ* = 2 and varying *J*_CF_ (left) and for fixed *J_CF_* = 0 and varying *κ* (right).

Next, we studied the migration of Hs578T cells in the computer simulations. Figure 5D shows the migration of the cells without cell-cell adhesion (*J*_CC_ = 0) for varying *κ* and *J*_CF_. Similar to the simulations for the 4T1 cells (with *J*_CC_ *>* 0), both cell-matrix adhesion and the traction force generated by the cells promote migration into the ECM. The measurement of the invasive radius confirms these observations (see Figure 5E).

Once again, we compared the intersection between the simulations and the experimental data to identify the mapping between the simulation parameters and the experiments with the knockouts. With the additional knockout in Hs578T cells, the computational model now proposes two possible scenarios of how the cell-matrix interactions change in the Hs578T cell line. For the Hs578T cells, Itg*α*2^-/-^cells can either 1) lose a significant amount of their cellmatrix binding but enhance their migratory capacity by generating more traction forces (intersection of data and simulations for *κ* = 3, *J*_CF_ = 0); or 2) lose part of their cell-matrix binding and moderately decrease their migratory capacity (*κ≈* 1.5, *J*_CF_ *>* 0). By performing additional qPCR tests (see Figure S4), we determined that the dimer *α*2*β*1 is the main integrin receptor expressed in these cells. But, as not all Itg*α*2 receptors were removed from Hs578T Itg*α*2^-/-^cells during experiments (see Figure 3A), this implies that cells only lose part of their cell-matrix binding and can still generate forces by pulling on the ECM during migration (see the black arrow in Figure 5E). Similarly, we can argue that Hs578T Itg*β*1^-/-^cells completely lose their cell-matrix binding, but still generate forces to moderately migrate by obtaining an ameboid migration mode. Thus, scenario 2 is the most plausible.

In addition to differences in invasiveness, the experiments also suggest that the knockouts change the morphologies of the invading cells. We characterize the shapes of the invasive cells in the simulations and show the shape distributions in Figure 5F. We note that these curves are consistent with earlier numerical work on cell shape distributions [56]. While the shape of the individual cells is hardly affected by the traction force generated by the cells (right panel), there is a clear shift towards more elongated cell shapes as the cell-matrix adhesion increases (left panel). In the absence of any cell-matrix binding, the cells exhibit a more round morphology. The experiments with Hs578T cells show more elongated cell morphologies when cell-matrix binding is present (sg control and Itg*α*2^-/-^), while the Itg*β*1^-/-^cells that detached from the tumoroid exhibit a more rounded morphology (see Figure 5A). These observations confirm that Itg*β*1^-/-^cells have no cell-matrix adhesion (*J*_CF_ = 0), while sg control and Itg*α*2^-/-^cells adhere to the ECM.

Overall, we can conclude that the impaired collagen binding introduced in the Hs578T and 4T1 cell lines mainly changes cell-matrix binding. The traction force generated per ECM binding site (described by *κ*) remains approximately the same for all the experiments. In the upcoming sections, we therefore focus on the computational results for a fixed value of the force generated per ECM binding site (*κ* = 2) and vary the cell-matrix adhesion strength *J*_CF_ together with the cell-cell adhesion strength *J*_CC_.

### C. Cell-cell adhesion enhances the formation of multicellular clusters

Since metastasis is ultimately seeded by detached cancer cell clusters [57–59], we next explored the properties of the cell clusters after they had separated from the primary tumor. In general, the detection and characterization of such clusters is difficult to achieve experimentally, but is relatively straightforward to measure in simulations. We thus studied the number of detached clusters and the size distribution of the invading clusters in simulations as a function of *J*_CC_ and *J*_CF_.

Figure 6A shows the total number of detachments as a function of *J*_CC_, for various values of *J*_CF_. Stronger cell-cell adhesion (larger *J*_CC_) leads to fewer detachments. These results are consistent with our intuition, as stronger cell-cell adhesion tends to suppress the breaking of contacts between neighboring cells. By contrast, an increase in the cell-matrix strength *J*_CF_ leads to more detachments. This can be understood by considering that cells adhere to the ECM and thereby migrate outward within our model. This outward migration leads to detachment from the primary tumor.

**Figure 6.**
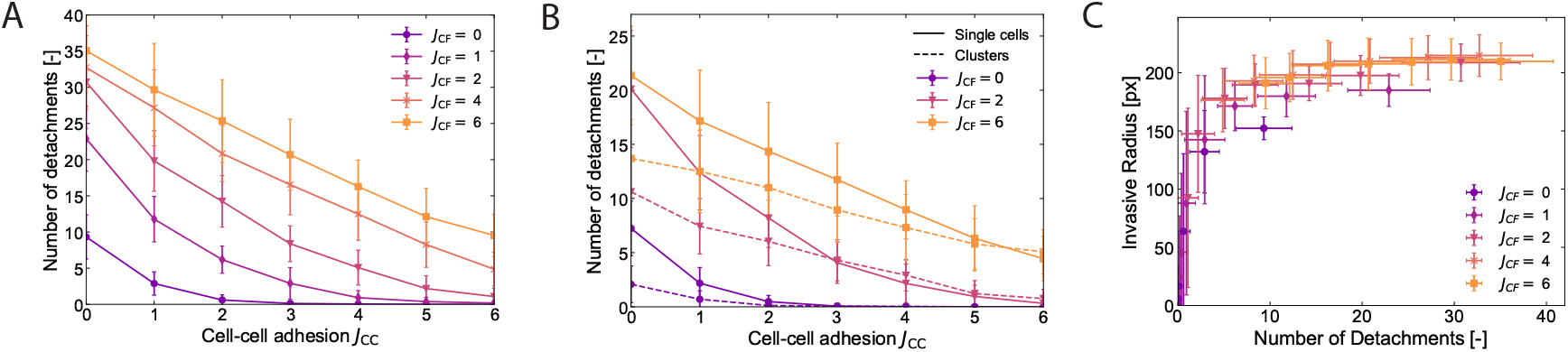
Number of detachments depends on the interplay between cell-cell and cell-matrix adhesion. **A**. Number of detachments as a function of the cell-cell adhesion strength *J*_CC_. The force generated by the invasive cells is kept constant at *κ* = 2. **B**. Number of single cells and multicellular clusters as a function of the cell-cell adhesion strength *J*_CC_ for the data shown in panel A (*κ* = 2). The data for *J*_CF_ = 1, 4 are left out for clarity. **C**. Correlation between the invasive radius and the number of detachments. *κ* is kept at a fixed value of 2.

Single detached cells are detected more frequently in our simulations than multicellular clusters (see Figure 6B), especially for low cell-cell adhesion strength. For high cell-cell adhesion strength, however, we detected an approximately equal number of single cells and multicellular clusters. Thus, within our model, stronger cell-cell adhesion promotes collective invasion. We note that cluster formation and collective migration can also be promoted by explicit cell-cell alignment or other forms of coordinated motion [60, 61], but in our model cell-cell adhesion is sufficient. When further discriminating between the different sizes of the multicellular clusters (see size distributions in Figure S8), we find that the probability of finding a cluster of a given size tends to decrease monotonically with size. These observed cluster size distributions are consistent with experimental observations [57] and other computational work [28, 62, 63]. Figure S8 also confirms that a higher cell-cell adhesion strength decreases the probability of single-cell detachments, in favor of multicellular cluster detachments.

Finally, we investigate how the number of detachments is related to the invasive radius. Increasing the cell-matrix adhesion (*J*_CF_) leads to both an increase in the invasive radius and the number of detachments, while a decrease in cell-cell adhesion (*J*_CC_) results in a decrease in both quantities. Indeed, Figure 6C shows that the invasive radius and the number of detachments are closely correlated, indicating that understanding one provides insights into the other.

Both quantities can serve as good proxies for describing the invasiveness of the primary tumor.

### D. Primary tumor morphology predicts invasiveness

Based on the previous sections, we investigate whether there are any other fundamental correlations that emerge as a predictive tool for dissemination. For example, the experimental data showed that the number of branches was reduced to a minimum when the tumor did not invade (see Figures 4 and 5), while a large number of branches may suggest active invasion, which could reflect increased cancer cell motility. Here, we assess the predictive value of our simulations by correlating the tumor morphology with its invasiveness for different cell-cell and cell-matrix adhesion strengths. While some of these predictions can be directly validated in experiments, the simulation model also offers predictions that go beyond our current experimental capability.

Figure 7A shows the correlation between the number of branches and the invasive radius measured in the simulations. The presence of a large number of branches is indicative of an invasive tumor, which is characterized by a high cell-matrix adhesion strength and a low cell-cell adhesion strength. The simulations suggest a non-linear relationship between the number of branches and the invasive radius. Most interestingly, for a given (fixed) value of the traction force per ECM binding site *κ*, we found that the data points for the different cell-cell and cell-matrix adhesion values collapse onto a single curve (see Figure S9 for results with different values of *κ*). This finding indicates that the invasive radius can be determined directly based on the number of branches, independent of the underlying values of the cell-cell and cell-matrix adhesion strength.

**Figure 7.**
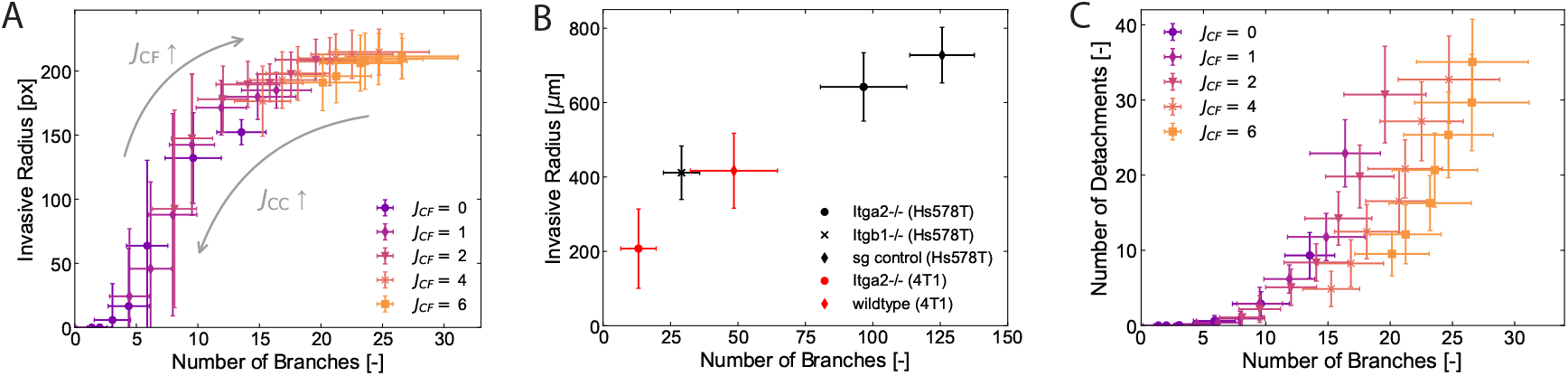
Data collapse of invasive radius versus number of branches, confirmed by experiments. **A**. Invasive radius as a function of the number of branches for the simulations with varying *J*_CC_ and *J*_CF_ (*κ* = 2) and **B**. Same results as in A for the experiments with Hs578T and 4T1 cells. Despite the different length scales between the simulations and experiments, these results agree qualitatively. **C**. The number of detachments correlated with the number of branches (*κ* = 2).

To confirm whether this prediction agrees with the experiments, we show the same data for the two cell lines in Figure 7B. Qualitative similarities between simulations and experiments can be observed. The experimental conditions in which the tumor is not invading (Hs578T-Itg*β*1^-/-^and 4T1-Itg*α*2^-/-^) are characterized by a small number of branches. On the other hand, the cells that can bind to the extracellular matrix create more branches and are more invasive, consistent with the simulation results. The results of the Hs578T and 4T1 cell lines also seem to lie on the same curve, suggesting that differences in cell-cell adhesion do not have a significant effect. Despite the differences in length scales, the invasive radius can thus be predicted based on the number of branches of the primary tumor core.

To further examine the predictive power of our computational model, we have also investigated other emerging correlations between morphology and invasiveness. An interesting candidate that could be used as a predictive marker is the complexity of the primary tumor, a measure for its shape. Since this marker is dimensionless, it has the additional benefit of being independent of the size of the primary tumor (see Figure S10 for collapse of data for different tumor sizes), which is not the case for the number of branches. Figure S11 shows the correlation between the complexity and the invasive radius for both the simulations and the experiments. The simulation results show that, similar to the number of branches, there is a direct correlation between the complexity of the primary tumor core and its invasive radius. Experimentally, the complexity was minimal when the tumor did not invade and the highest complexity was observed in the wildtype cells that invaded the most (see Figure S5). However, the correlation that is predicted by the simulations is not apparent in the experimental data (see Figure S11). We explain this discrepancy by the fact that the complexities for wildtype 4T1 and Hs578T cells were very similar, while the morphologies of the primary tumor were different. Similarly, the complexities of the Hs578T wildtype and Itg*α*2^-/-^cells were almost identical while the invasive radius was significantly reduced. Further characterization of the shape of the primary tumor would be necessary to make an accurate prediction about its invasive radius.

Finally, we investigate the relationship between the number of branches and the number of detachments. Figure 7C shows that while there is a correlation, it is more spread compared to previous measures. This indicates that predicting the number of detachments directly from the number of branches is more challenging. Interestingly, as the number of detachments remains the same, we observe that increasing cell-matrix adhesion (*J*_CF_) leads to more branches. Based on the experimental data, the correlation that we find has some merit (see Figure S12), but it must be noted that extracting the number of detachments in the experiments reliably between the different biological replicates is challenging. Overall, this nuanced relationship between the number of detachments and the number of branches underscores the difficult interplay of cell-cell and cell-matrix interactions in tumor invasion dynamics.

### E. Threshold of invasion depends linearly on cell-cell adhesion and cell-matrix interactions

Until now, we have focused on the role of cell-cell and cell-matrix adhesion on the invasion of cells. In this final result section, we go beyond the experimentally accessible conditions and focus on the strength of the traction force parameter *κ*. This parameter could not be varied with the two cell lines and integrin knockout experiments, but our computational model allows us to study how traction forces affect cell invasion.

Figures 4C and 5E have already shown that increasing *κ* leads to an increase in the invasive radius and thus promotes invasion of cells into the ECM.

More interestingly, for strong cell-cell adhesion, the model suggests that the cells need to generate a certain amount of traction forces before invasion occurs (see Figure 4C around *κ* = 2). This so-called invasion threshold depends both on the cell-cell adhesion and the cell-matrix interactions. To map out the invasion regimes, we explored the full parameter regime (described by *J*_CC_, *J*_CF_ and *κ*) and identified the threshold for invasion. Here we define this threshold as the point where the mean invasive radius exceeds the original size of the primary tumor core (equal to 100 pixels). Other measures, such as determining the number of simulations in which cells detached from the primary tumor, led to slight changes in the onset of invasion (data not shown), but the general trends remained the same.

Figure 8A shows the invasive radius as a function of cell-cell adhesion strength *J*_CC_ and traction force per ECM binding site *κ* for *J*_CF_ = 1. For strong cell-cell adhesion strengths (*J*_CC_ *>* 4) and small traction forces (*κ* = 1), the primary tumor is noninvasive, whereas a lack of cell-cell adhesion (*J*_CC_ = 0) leads to migration into the ECM. The threshold for invasion (gray line) suggests a quasi-linear relationship between the strength of the cell-cell adhesion and the traction force generated by the individual cells. This linear relationship between *J*_CC_ and *κ* also holds for other values of the cell-matrix adhesion strength *J*_CF_ (see Figure 8B).

**Figure 8.**
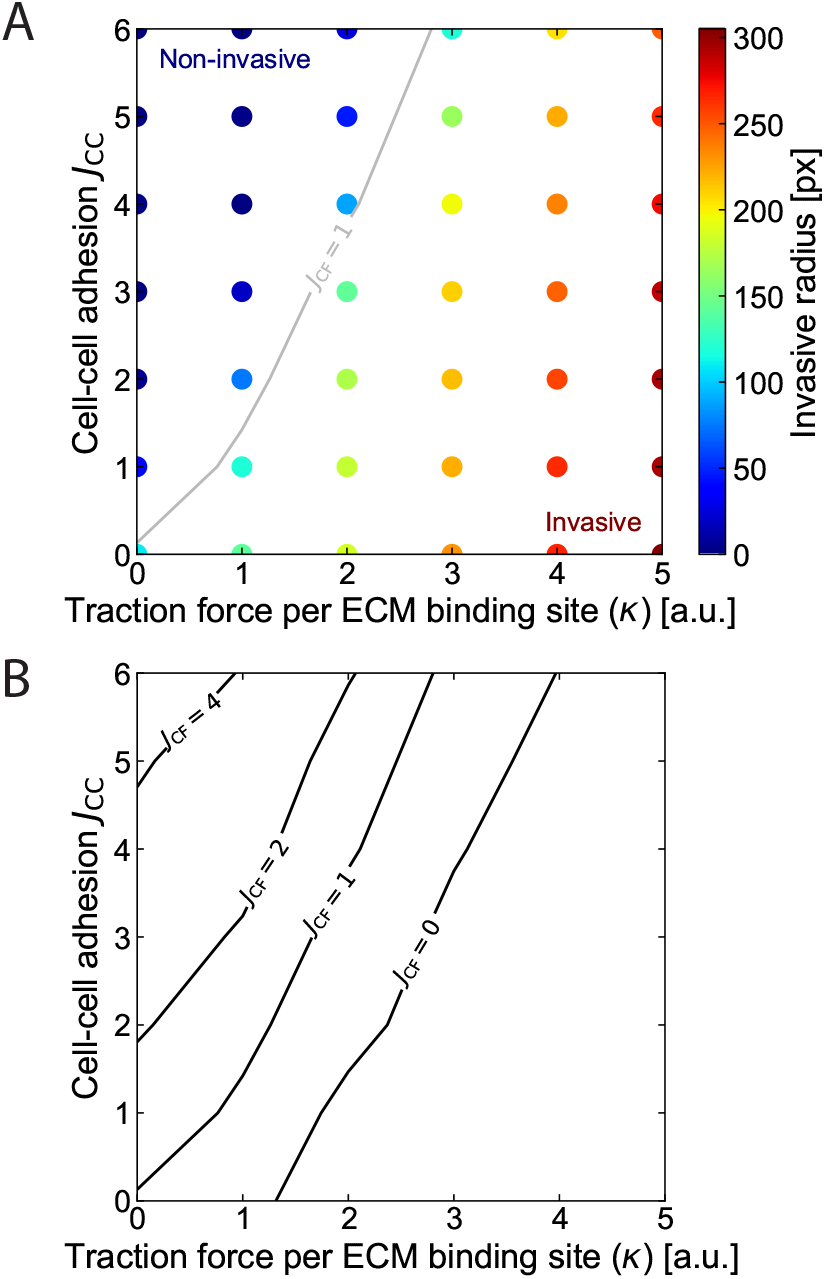
State diagram with the onset of invasion as a function of cell-cell and cell-matrix interactions. **A**. Mean invasive radius as a function of *κ* and *J*_CC_ for *J*_CF_ = 1. The threshold for invasion is indicated in gray. **B**. Threshold for invasion shifts towards stronger cell-cell adhesion strength and smaller traction forces for increasing *J*_CF_ since cell-matrix adhesion promotes invasion.

To interpret the observed invasion threshold in the context of physical adhesion models, we draw an analogy with the Johnson-Kendall-Roberts (JKR) model [64]. In this model, the adhesion between two elastic surfaces scales linearly with the pull-off force required to separate them. Similarly, in our simulations, the traction force parameter *κ* can be viewed as the cellular equivalent of this pull-off force, counteracting the adhesive interactions between cells. The quasi-linear relationship we observe between the required traction force and the strength of cell-cell adhesion is similar to the JKR prediction, suggesting that the onset of invasion may be governed by a local force balance at the single-cell level. Notably, this linear relationship persists across different values of the cell-matrix adhesion strength *J*_CF_. This may be explained by the fact that increasing *J*_CF_ effectively increases the number of ECM binding sites, which in turn leads to a proportional increase in the traction force. The analogy with the JKR model implies that invasion is not merely a collective property of the tumor but may emerge from the interaction between cell-cell adhesion and traction forces on the scale of individual cells.

## 4. DISCUSSION AND CONCLUSION

In this study, we investigated the role of cell-cell and cell-matrix interactions in cancer cell invasion by combining integrin knockout experiments in Hs578T and 4T1 cell lines with a computational cellular Potts model. Our results show that the invasive behavior of these cell lines can be captured by varying cell-cell adhesion parameters in the model, with Hs578T and 4T1 cells corresponding to low and high cell-cell adhesion regimes, respectively. We further demonstrate that integrin knockouts (Itg*α*_2_ and Itg*β*_1_) can be modeled by reducing cell-matrix adhesion strength (*J*_CF_), while maintaining a consistent traction force per ECM binding site (*κ*). These findings establish the computational model’s capacity to replicate key features of the experimental invasion dynamics, supporting its utility for predictive modeling.

As part of our predictive modeling approach, we show that tumors with highly irregular shapes, characterized by a large number of branches, exhibit greater invasiveness, increasing the likelihood of cell dissemination. Such correlations between tumor morphology and invasive potential are difficult to extract experimentally because of the limited ability to systematically vary biological parameters across a wide range of conditions. Our computational model enables controlled exploration of cell-cell and cellmatrix adhesion strengths, allowing us to map the invasive radius as a function of morphological features. In particular, for a given *κ*, the data collapses onto a single curve when plotting the invasive radius against the number of branches, suggesting that detailed knowledge of the adhesion parameters may not be required to estimate the invasive radius. This finding points to the potential of using tumor shape as a proxy for invasiveness, even in the absence of detailed cellular information. While promising, further work is needed to assess whether other features, such as the number of cell detachments, can also be inferred from morphological descriptors.

Finally, we show that traction forces play a critical role in the onset of invasion. Measurement of these forces in 3D environments remains experimentally challenging, as it requires embedding of soft deformable microparticles within the ECM and tumoroid, which is very time-consuming and labor-intensive [65–67]. Computational models therefore provide a valuable tool for gaining mechanistic insight into how traction contributes to invasive behavior. Previous studies have established that active motility, similar to our traction force, is essential for invasion [50, 51], and increased traction has been correlated with enhanced invasiveness in experimental systems [52, 68], consistent with our findings. Our results suggest that the onset of invasion is governed by a local balance between cell-cell and cell-matrix interactions. Further characterization of this invasive regime – such as distinguishing whether cells invade as isolated single cells, as coordinated multicellular groups, or remain non-invasive [25, 33] – may help clarify how these interactions determine the specific mode of invasion.

While our model is developed in close alignment with experimental observations, it is important to recognize the assumptions that influence its predictions, particularly regarding the role of cell-matrix adhesion. Our simulations indicate that increasing cell-matrix adhesion promotes invasion, consistent with previous computational studies [31, 51]. However, other studies have reported the opposite effect. For example, Steijn *et al*. [37] found that strong adhesion can hinder invasion by anchoring cells to ECM binding sites and reducing their motility. This difference can be explained by the underlying assumptions of our model: we assume that the traction force scales with the number of ECM binding sites, so stronger adhesion enhances motility [52]. If this assumption is removed by decoupling the traction from the number of the binding site, our model should reproduce the pinning effect described by Steijn *et al*. [37], where stronger adhesion limits invasion. These results highlight how the relationship between adhesion and invasion depends on the specific mechanistic coupling between adhesion and force generation.

One limitation of our computational model is its use of static ECM fibers to represent the collagen network observed in experiments. In reality, the extracellular matrix plays an active role in tumor progression, as cancer cells dynamically remodel the ECM to facilitate invasion and metastasis [69]. A common feature of invasive tumors is the radial alignment of ECM fibers around the tumoroid, known as the tumor-associated collagen signature (TACS) [70–72]. This radial organization enhances invasion by providing directional guidance for migrating cells [73–75]. Consistent with previous modeling studies [44, 51], our results show that ECM topology strongly influences migration: a radially aligned ECM promotes outward movement and increases invasiveness. Incorporating ECM remodeling into future versions of the model could provide a more realistic representation of the dynamic interplay between cells and their microenvironment.

Another natural extension of this work involves enhancing the model to better capture the complexity of large-scale collective invasion. While we observe the emergence of small multicellular strands, the model does not fully reproduce the robust, extended strand formation reported in experimental studies [27, 34, 63]. Prior modeling work has shown that cell-cell adhesion alone, even when combined with an active leading cell, is insufficient to drive sustained strand formation over long distances [76]. This points to the need for additional mechanisms – such as leader-follower interactions – that may be necessary to coordinate and sustain collective invasion [77–79]. In particular, the emergence of cellular heterogeneity, where distinct subpopulations adopt specialized migratory roles, may play a key role in enabling large-scale strand formation. Incorporating such heterogeneity into future versions of the model, for example through explicit representation of leader-follower behavior, may improve the model’s capacity to capture the full range of collective invasion behaviors.

## ACKNOWLEDGMENTS

The authors thank Rick Rodrigues de Mercado for his help with adapting the MATLAB code for tumoroid analyses, Leon Hillmann for developing the shape analysis code, and Hanneke van de Sanden for help with preliminary simulation studies. This project is funded by the Dutch Research Council (NWO) through the ENW-XL project “Active Matter Physics of Collective Metastasis” (OCENW.GROOT.2019.022).

