## Supplemental Materials for "Adhesion-driven invasion: disentangling the interplay between cell-cell and cell-matrix interactions in cancer cell migration"

### SUPPLEMENTAL MATERIAL

#### S1. Computational Model

To investigate the interplay between cell-cell and cell-matrix interactions in cell migration, we developed a computational model based on the cellular Potts (CP) formalism [39, 40]. Briefly, our model consists of a static, fibrous ECM structure and a dynamic primary tumor core (see Figure 1A). In reality, cells experience complex interactions with the ECM in three dimensions (3D), including squeezing through pores and navigating complex tissue architectures. However, we opted for a quasi-two-dimensional setup for computational simplicity, as it also allows us to study the competition between cell-cell adhesion and different cell-matrix interactions during migration in a tractable manner. Our simulations thus represent a plane section of the tumoroid. The model is implemented in CompuCell3D [41].

Within our setup, the ECM fibers function as a guide to the tumor cells through cell-matrix interactions. We represent the ECM fibers as discrete fibers on a square lattice, similar to previous modeling work [30, 35, 42–45]. The orientation of these fibers is based on experimentally observed ECM networks (see Figure 1C). In particular, cells actively remodel the ECM during cell migration, causing initially randomly-oriented collagen fibers to reorient in a radially outward direction [69–72]. Here we consider the two limiting cases of this remodeling process, i.e. a random ECM and a radial ECM topology. Since we are mainly interested in cell migration and invasion into the ECM, we focus primarily on results obtained with the radial ECM topology; The random ECM topology is used as a benchmark in absence of any ECM remodeling.

The tumoroid is placed on top of the ECM fiber configuration (see Figure 1A). The primary tumor core contains two distinct cell types: nonmotile tumor cells and motile invasive cells. Both cell types possess the same specific volume and surface properties. The surface term reflects the membrane of a cell and its cortical tension, while the volume term accounts for the compressibility of the cell [46]. In addition to these intracellular cell properties, the cells can adhere to other cells and to ECM fibers. We denote the cell-cell adhesion strength with $J_{\mathrm{CC}}$ and the cell-matrix adhesion strength with $J_{\mathrm{CF}}$. In the absence of either cell-cell adhesion or cell-matrix adhesion, we set the adhesion strengths equal to zero. When cells prefer to adhere to the ECM fibers, $J_{\mathrm{CF}}$ increases to promote cell-matrix binding. An increase in $J_{\mathrm{CC}}$, on the other hand, promotes cell-cell contacts such that cells are more strongly bound to each other. Thus, by adjusting the values of $J_{\mathrm{CC}}$ and $J_{\mathrm{CF}}$, we can modify the adhesion interactions with the extracellular matrix and other cells.

The motile invasive cells also interact with the ECM through force generation, allowing them to actively migrate outwards. Here we define invasive cells as those that form the boundary of the primary tumor and that are in contact with at least one ECM fiber (see zoom-in in Figure 1A). The traction force $\vec{F}_a(t)$ generated by the invasive cells due to contact with the ECM is modeled as

$$\vec{F}_a(t) = \kappa N_{\mathrm{CF}}(t)\hat{p}_\sigma(t), \tag{2}$$

where $N_{\mathrm{CF}}(t)$ is the number of ECM contacts at a given time $t$, $\hat{p}_\sigma(t)$ is the polarity vector that determines the direction of the force, and the scaling factor $\kappa$ is the force generated per ECM binding site. For the time evolution of $\hat{p}_\sigma(t)$, we follow previous works [48–51] and impose a self-alignment mechanism in which the cell aligns its polarity with its previous velocity direction [47]. This ensures that the invasive cells migrate radially outwards into the ECM. Note that the traction force scales linearly with the number of ECM contacts. Experiments have shown that an increase in the collagen density can lead to an increase in the traction forces generated by the cells [52]. Although our linear dependence is arguably the simplest implementation, we have verified that a parabolic dependence on $N_{\mathrm{CF}}$ leads to the same phenomenological behavior. An overview of the cell-cell and cell-matrix interactions that are included in the computational model are schematically shown in Figure 1B.

We run the simulations with an initial primary tumor core of 316 cells and an initial radius of approximately 100 pixels. The ECM contains 5000 ECM fibers, with an end-to-end length of 10 pixels each, such that the average area covered by the ECM fibers is around 17%. For each simulation, we generate a new ECM fiber network, with the centers-of-mass of the fibers positioned randomly in space. The orientation of the fibers is either random (random ECM) or radially outward (radial ECM).The initial configurations of the cells are equilibrated for 1000 Monte Carlo steps (mcs) without any cell interactions ($\kappa = 0, J_{\mathrm{CC}} = 0, J_{\mathrm{CF}} = 0$), after which the simulations are run for another 5000 mcs including the cell interactions. During these 5000 mcs, the most invasive tumors grow to approximately 3 to 4 times their original size. We studied the migratory cell behavior as a function of the scaling of the traction force ($\kappa$), the cell-cell adhesion strength ($J_{\mathrm{CC}}$), and the cell-matrix adhesion strength ($J_{\mathrm{CF}}$). An overview of the complete set of simulation parameters, including the definition of the Hamiltonian in the CP formalism, is provided in the Supplementary Material.

### S2. Details of the Cellular Potts Model

The dynamics in the Cellular Potts (CP) model is drived by the following Hamiltonian,

$$\mathcal{H} = \sum_{k,l \in \text{neighbors}} -J_{\tau_k,\tau_l}(1-\delta_{kl}) + \sum_i \left[\lambda_V (V_i - V_T)^2 + \lambda_S (S_i - S_t)^2\right] - \sum_i \kappa N_{CF,i} \hat{p}_i \cdot \vec{R}.$$

The index $i$ indicates the cell number, while the indices $k, l$ refer to specific lattice sites. The cell type is defined by $\tau_k$. There are three main contributions to the Hamiltonian:

- Adhesion contributions: cells can adhere to the different components in the simulations. The cell-cell adhesion ($J_{\mathrm{CC}}$) is the adhesion strength between the cells that are either of type 'invasive' cell or 'tumor' cell. Similarly, these cells can adhere to the ECM fibers with an adhesion strength of $J_{\mathrm{CF}}$. By default, the values are equal to 0 (no adhesion) and increasing the value for adhesion ($J_{\tau_k,\tau_l} > 0$) increases the adhesion strength.

- Volume and surface constraint: cells have a preferred volume ($V_T$) and a preferred surface ($S_T$). These are enforced by the second term in the Hamiltonian. The strength of the constraint is tuned by $\lambda_V$ and $\lambda_S$.

- Cell motility: the cells can generate forces to migrate. Only invasive cells can generate forces in response to the ECM fibers. The number of contacts a cell has with an ECM fiber in the underlying substrate is $N_{\mathrm{CF}}$. The strength of this force is scaled with a factor $\kappa$. The direction of migration is determined by the velocity direction of a cell.

Based on the Hamiltonian, the dynamics is driven by the modified Metropolis algorithm [80],

$$P(\tau_k \to \tau_{k'}) = \begin{cases} 1 & \Delta\mathcal{H} \leq 0 \\ \exp\left(-\frac{\Delta\mathcal{H}}{T_m}\right) & \Delta\mathcal{H} > 0, \end{cases}$$

where $T_m$ is an effective temperature that determines the membrane fluctuations. The full set of parameters that is used in the simulations is provided in Table S1.

| Parameters | Values | Description |
|---|---|---|
| $T_m$ | 10.0 | Temperature (membrane fluctuations [41, 46]) |
| Box size | (600 px, 600 px, 2) | (x, y, z) coordinates |
| $\lambda_V$ | 10.0 | Strength of the volume constraint |
| $V_T$ | 100.0 px | Preferred cell volume |
| $\lambda_S$ | 0.25 | Strength of the surface constraint |
| $S_T$ | 135.0 px | Preferred cell perimeter |
| $J_{\mathrm{CM}}$ | 0.0 | Adhesion between Cells and Medium |
| $J_{\mathrm{CC}}$ | [0; 1; 2; 3; 4; 5; 6] | Adhesion between Cells |
| $J_{\mathrm{CF}}$ | [0; 1; 2; 4; 6] | Adhesion between Cells and Fibers |
| $\kappa$ | [0; 1; 2; 3; 4; 5] | Scaling factor for the force generated by an invasive cell |
| $N_{\mathrm{F}}$ | 5000 | Number of fibers |
| $L$ | 10.0 px | End-to-end length of the fibers |
| $t_{\mathrm{eq}}$ | 1000 mcs | Equilibration time |

Table S1. Complete set of simulation parameters. The parameter $J_{\mathrm{CC}}$, $J_{\mathrm{CF}}$ and $\kappa$ are varied in the results section of the main text.

### S3. Computational artifacts in the large force regime

In the computational model, we focus on the invasive radius as a measure to characterize the migration of the cells. In the end, we aim to find a correlation between the invasive properties and the shape of the primary tumor core, which can potentially be used as a predictor for the overall invasiveness of a tumor. We use the number of branches as a measure for the shape of the primary tumor core. We also studied if the complexity could be used as a predictive marker for invasion. The complexity can, however, only be used accurately in the regime where the traction forces generated by the cells are small. For cells that strongly bind to the ECM and thereby generate large traction forces, the primary tumor core disintegrate in many small clusters as the cells are strongly invasive. In this regime, the primary tumor core becomes ill-defined and the complexity actually decreases. In this regime, the complexity is no longer a meaningful predictor of the invasiveness of the primary tumor core. Therefore, we only use the results for $\kappa \leq 2$ in Section 3 D.

### S4. Experimental methods

#### Cell culture

Cell lines Hs578T and 4T1 were obtained from the American Type Culture Collection and cultured in high-glucose Dulbecco's Modified Eagle's Medium (DMEM, Gibco, 11504496, MA, USA) containing L-Glutamine and Sodium Pyruvate, supplemented with 10% fetal calf serum (Thermo Scientific, MA, USA) and 25 μg/mL penicillin/streptomycin (Gibco, 15070-063, MA, USA) in a humidified incubator at 37°C with 5% $CO_2$.

**RT-qPCR**

For RT-qPCR, total RNA was isolated by RNeasy Plus Mini Kit (Qiagen, 74134, Germany) and cDNA was synthesized by the RevertAid H Minus First Strand cDNA Synthesis Kit (Thermo Fisher Scientific, K1632, MA, USA). Real-time qPCR was performed with 2 biological replicates in quadruplicates using PowerUp SYBR Green Master Mix (Thermo Fisher Scientific, A25742, MA, USA) on QuantStudioTM 6 Flex Real-Time PCR system (Applied Biosystems). The qPCR primer sets can be found in the Supplementary Material. Relative mRNA expression was calculated and normalized to the control (ACTB) using the $2{-}\Delta CT$ method. The following mouse qPCR primer sets were used:
ACTB forward (fw), 5'-CTAAGGCCAACCGTGAAAAG-3';
ACTB reverse (rev), 5'-ACCAGAGGCATACAGGGAC-3';
ITGB1 forward (fw), 5'-CGGACGCTGCGAAAAGATGA-3';
ITGB1 reverse (rev), 5'-CTTGGCTGGCAACCCTTCTT-3';
ITGA2 forward (fw), 5'-AGGACCTTTGCTGCTTCGAC-3';
ITGA2 reverse (rev), 5'-TGCTTTCTCCGTGGGTTTCA-3';
ITGA1 forward (fw), 5'-ATGACGCTCTGCCAAACTCA-3';
ITGA1 reverse (rev), 5'-TGTTGTACGCACTGTCTCCC-3';
ITGA10 forward (fw), 5'-CCAGGATTCTCGGTTTGGCT-3';
ITGA10 reverse (rev), 5'-GAAGTATCGGAGGGCCTGTG-3';
ITGA11 forward (fw), 5'-AGCCTTTGGCCAGGATTCAC-3';
ITGA11 reverse (rev), 5'-AGGTTGAGCTTGGTGCAGTT-3';
DDR1 forward (fw), 5'-CTTTGGCTTCAGGTGCCTCG-3';
DDR1 reverse (rev), 5'-GGGGACAGCATCTCTCCAGG-3';
DDR2 forward (fw), 5'-CACCCACCACCTATGATCCC-3';
DDR2 reverse (rev), 5'-CTTGGTGATGAGGAGCGGTT-3';
CDH1 forward (fw), 5'-GGCTGGACCGAGAGAGTTAC-3';
CDH1 reverse (rev), 5'-CCGGGCATTGACCTCATTCT-3';
CDH2 forward (fw), 5'-GGCCTTGCTTCAGGCGTC-3';
CDH2 reverse (rev), 5'-AGGAACTTTGCCTGCTCTGC-3';
The following human qPCR primer sets were used:
ACTB forward (fw), 5'-CTTCGTGTCCTGTATGGCCC-3';
ACTB reverse (rev), 5'-ACAGAGGTAGGTGCCCTCAA-3';
ITGB1 forward (fw), 5'-GCCGCGCGGAAAAGATGAA-3';
ITGB1 reverse (rev), 5'-CCACCCACAATTTGGCCCTG-3';
ITGA2 forward (fw), 5'-CGGTTATTCAGGCTCACCGA-3';
ITGA2 reverse (rev), 5'-GCTGACCCAAAATGCCCTCT-3';
ITGA1 forward (fw), 5'-GAAATCACTCCTCTCCCGCT-3';
ITGA1 reverse (rev), 5'-AGCAGCGTAGAACAACAGTG-3';
ITGA10 forward (fw), 5'-GACGTTTATCGCTGCCCTGT-3';
ITGA10 reverse (rev), 5'-CCTGAGGCTGGAATGAAGCA-3';
ITGA11 forward (fw), 5'-CCGCTCTGGCTTGCCG-3';
ITGA11 reverse (rev), 5'-GGTGTCCGTGAACCCTGG-3';
DDR1 forward (fw), 5'-GCCACCAAGAATGCCAGGAA-3';
DDR1 reverse (rev), 5'-CAGCAGCATTGGGTAGCTGA-3';

DDR2 forward (fw), 5'-CTCCTTCACCAGCAAAGTGGA-3';
DDR2 reverse (rev), 5'-TCCCTTGATGGAGGTTTCATCC-3';
CDH1 forward (fw), 5'-TCGCTTACACCATCCTCAGC-3';
CDH1 reverse (rev), 5'-CCTGACCCTTGTACGTGGTG-3';
CDH2 forward (fw), 5'-GGACAGCCTCTTCTCAATGTG-3';
CDH2 reverse (rev), 5'-CTTCTGCTGACTCCTTCACTG-3';

**Lentiviral genomic modifications**

For stable 4T1 knockout cells, a lentiCRISPR-Cas9 vector (Addgene, 52961, MA, USA) was digested using BsmBI (New England Biolabs, R0739S, MA, USA), FastDigest buffer (Thermo Fisher Scientific, B64, MA, USA), FastAP Thermosensitive Alkaline Phosphatase (Thermo Fisher, EF0654, MA, USA) and 1 mM DTT (Intvitrogen, Y00147, MA, USA). DNA of digested plasmid was separated on a 0.8% agarose gel and a 13 kbp band was cut out and purified using Wizard SV Gel and PCR Clean-Up System (Promega, A9282, WI, USA). Three guide RNAs were designed for ITGA2: TGGCCAGTATAATTTGCTCG, TACAACTACAACATCCACGA and TCAGAGCTATCGAATTCGCA. Three guide RNAs were designed for ITGB1: GAGGAATGTAACACGACTGC, TCCCAACATTCCTACCAATG and GGAAATGGGACATTTGAGTG. Oligos for each guide RNA were ordered with caccg-overhang on the 5'-end of the forward, and c-overhang on the 3'-end and caaa-overhang on the 5'-end of the reverse oligo (Sigma, MO, USA). Each pair of oligos was phosphorylated and annealed with T4 DNA Ligase Reaction Buffer (New England Biolabs, B0202S, MA, USA) and T4 PNK Ligase (New England Biolabs, M0201S, MA, USA) in a thermocycler (Mastercycler Nexus gradient, Eppendorf, 63331, Germany) at 37°C for 30 minutes and 95°C for 5 minutes. Oligos were ligated with digested plasmid with Quick Ligation kit including 2X Quick Ligase Buffer and Quick Ligase (New England Biolab, M2200S, MA, USA) and incubation at room temperature. Ligated plasmids were transformed into Stbl3 E.coli bacteria (Thermo Fisher Scientific, C737303, MA, USA) and colonies were picked and expanded. DNA was isolated using a Plasmid Midiprep kit (LabNed, LIV2400004, The Netherlands) and sanger sequencing with a primer binding to the LKO1 sequence (gactatcatatgcttaccgt) was performed to validate the cloning.

For doxycycline inducible Hs578T Cas9 cells an Edit-R Inducible Lentiviral hEF1$\alpha$-Blast-Cas9 Nuclease Plasmid DNA (Dharmacon, CAS11229, CO, USA) was combined sequentially with U6-gRNA/PGK-Puro-2A-BFP vectors expressing short guide RNAs (sgRNAs) targeting ITGA2: CCCAGAAGCAAAAATATTTTCCG, CTGGTTGGTTCACCCTGGAGTGG; or ITGB1: CCAGTGTAGTTGGGGTTGCACTC, CCAAAGAACAGTCACATGCACTG, which were obtained from the Sanger Whole Genome CRISPR Library (Sigma-Aldrich). Additionally a non-targeting sgRNA was used as negative control. For virus production, a DNA mixture was prepared combining the third generation lentiviral vectors described above (either Edit-R Inducible Lentiviral hEF1$\alpha$-Blast-Cas9 or U6-gRNA/PGK-Puro-2A-BFP) with helper constructs pMDLg-RRE, pCMV-VSVG and pRSV-Rev (all from Addgene, 12251, 8454, 12253 respectively, MA, USA). Subconfluent Lenti-X 293T cells (Clontech, 632180, CA, USA) were transfected with the DNA mixture and Polyethylenimine (Polysciences, 23966-2, PA, USA). After 24 hours the medium was refreshed, after 48 hours and 72 hours the medium was collected and filtered through a 0.45 μm filter. For lentiviral transductions, 50,000 cells were adhered in a 6-wells plate (Greiner Bio-one, 657160, Austria) and virus containing medium was added and supplemented with 5 μg/mL Polybrene (Sigma-Aldrich,

MO, USA). After 24 hours the medium was changed. The inducible Hs578T-Cas9 cell line was selected using 2 μg/mL Blasticidin (Sigma-Aldrich, 203350, MO, USA) followed by isolation of a single clone that was expanded and tested for Cas9 expression upon Doxycycline (Selleckchem, S5159, TX, USA) exposure. For sgRNA U6-gRNA/PGK-Puro-2A-BFP plasmids, the following guide RNAs were added into U6-gRNA/PGK-Puro-2A-BFP vector by the LUMC: ITGA2: CCCAGAAGCAAAAATATTTTCCG, CTGGTTGGTTCAC-CCTGGAGTGG; ITGB1: CCAGTGTAGTTGGGGTTGCACTC, CCAAAGAACAGTCACATGCACTG. Additionally a non-targeting sgRNA was used as negative control.

After lentiviral transductions of three sgRNAs in 4T1 and two in Hs578T-Cas9, cells were selected using puromycin (1 μg/mL for Hs578T and 8 μg/mL for 4T1, Gibco, A1113803, MA, USA). Cas9 expression in Hs578T-Cas9 was induced with 1 μg/mL Doxycycline for a minimum of 7 days.

#### Western blot

Knockouts were verified with Western blot. Twenty μg of protein per sample were loaded on sodium dodecyl sulfate–polyacrylamide gels, with acrylamide concentrations ranging from 7.5% to 4%. Protein lanes were transferred to polyvinylidene difluoride membranes (Millipore, IPVH00010, MA, USA). Cas9 (Cell Signaling Technology, 14697, The Netherlands), Itgb1 (Millipore, AB1952, MA, USA) and Itga2 (Abcam, ab181548, United Kingdom) antibodies were used for stainings. Membranes were visualized with Amersham ECL Western Blotting Detection Kit (GE HealthCare, RPN2232, IL, USA). All uncut, original membranes used in the figures are shown in Figure S1.

#### Proliferation and adhesion

To determine the effect of knockout on proliferation, 4.000 cells were plated in 96-well plates (Gibco, 655180, ThermoFisher Scientific, MA, USA). Cell proliferation was assessed using the IncuCyte® S3 (Sartorius, Germany), equipped with a 10x objective and placed inside an incubator with 37°C and 5% $CO_2$. Images of live cells were acquired every 2 hours over 3 days in phase contrast channel. The images were masked and confluence was assessed using the Incucyte software version 2020B (Essen BioScience Inc.). Collagen type I solution was isolated from rat-tail collagen by acid extraction as described previously [81]. For collagen adhesion assay, 4.000 cells were plated in plates that were coated with 20 μg/mL rat tail collagen I mixed in water. All cell lines were plated at the same time and for 4T1, 30 minutes and for Hs578T, 10 minutes after incubation at 37°C, plates were washed twice to obtain the loss in binding to collagen. Then, plates were placed back into the incubator and incubated overnight. Cells were fixed with 2 % formaldehyde (Sigma-Aldrich, 252549, MO, USA) and stained with 0.1 % TritonX-100 (Sigma-Aldrich, T8787, MO, USA), 0.4 μg/mL Hoechst 33258 (Invitrogen, H21491, MA, USA) and 0.06 μM Phalloidin Rhodamine (ThermoFisher Scientific, R415, MA, USA) in PBS for 1 hour and then washed twice.

#### Tumoroid formation

Collagen matrices were created by mixing with DMEM (Gibco, ThermoFisher Scientific, MA, USA), HEPES 0.1 M (1M stock, Biosolve, The Netherlands) and $NaHCO_3$ 44 mM (Sigma-Aldrich, S8875, MO,

USA) as previously described [15]. 30 µl collagen mixture was polymerized in a 384-well CELLSTAR® plate (Greiner Bio-one, 781091, Austria) at 37°C for 1 hour.

Spheroid or Tumoroid formation into 3D collagen matrix was performed as described previously [82]. Subconfluent monolayers of tumor cells were trypsinized and filtered (Sysmex, 04-0042-2317, Japan). Cells were resuspended in PBS containing 2 % polyvinylpyrrolidone (PVP, Sigma-Aldrich, P5288, MO, USA) to reach a cell concentration of approximately 50.000 cells per µl. Immediately after, tumoroids, 200 µm in diameter and containing approximately 2.500 cells, were created by automated printing of cell-PVP mixture into the collagen-matrigel matrix at defined x-y-z positions 150 mm above the bottom of the wells using the injection robotics from Life Science Methods (Leiden, The Netherlands). After injection, tumoroids were incubated with cell culture medium at 37°C. After 24 hours tumoroids were fixed with 2 % formaldehyde and stained with 0.1 % TritonX-100, 0.4 µg/mL Hoechst 33258 and 0.06 µM Phalloidin Rhodamine in PBS for 3 hours at room temperature. The samples were washed thoroughly in PBS and microscopy was performed shortly after.

### Scanning confocal microscopy

Images of tumoroids were acquired on a Nikon Eclipse Ti inverted scanning confocal microscope equipped with four laser lines 405 nm, 488 nm, 561 nm and 640 nm, and with an A1R MP scanner. The microscope has a Nikon encoded and automated stage. Its camera is controlled through NIS Element Software (Nikon Instruments Inc., Melville, NY, USA). Tumoroids were imaged with 20 µm distance between Z-slices. ECM collagen-fibers were detected by confocal reflection microscopy on the same scanning confocal microscope. The collagen fibers were scanned at 561 nm excitation with a 561 nm blocking dichroic mirror and all the light reflected back passed a through filter of bandwidth 400 nm - 750 nm. The reflected light from the collagen fiber network was collected on GaAsP-photomultiplier. Scanning confocal fluorescent microscopy of tumoroids, and scanning confocal reflection microscopy of the collagen network were performed using Plan Apo ×20/0.75 NA, and Apo LWD ×20/0.95 objectives (Nikon Instruments Inc., Melville, NY, USA), respectively. Images of fixed tumoroids were acquired with laser lines 405, 488, 561, Transmission detection channel and reflection microscopy.

### Statistics

An unpaired, parametric t-test was used to determine statistical significance between populations. Data sets were significantly different with probabilities of $p < 0.0001$ (****), $p < 0.001$ (***), and not significantly different for probabilities of $p > 0.05$ (ns).

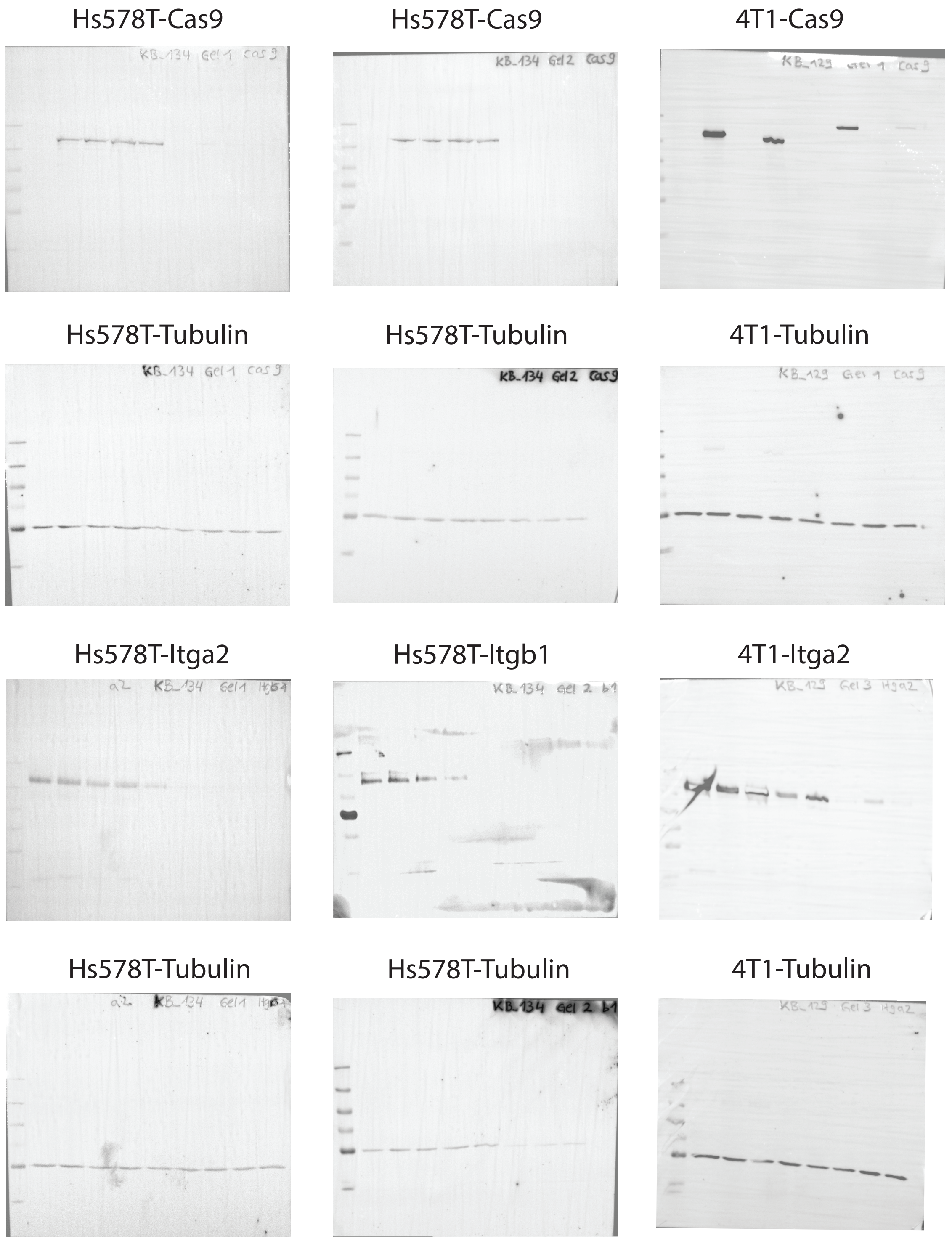

Figure S1. **Uncut Western blot membranes.** Stainings show complete integrin knockouts in Hs578T and 4T1 cells according to Figure 3.

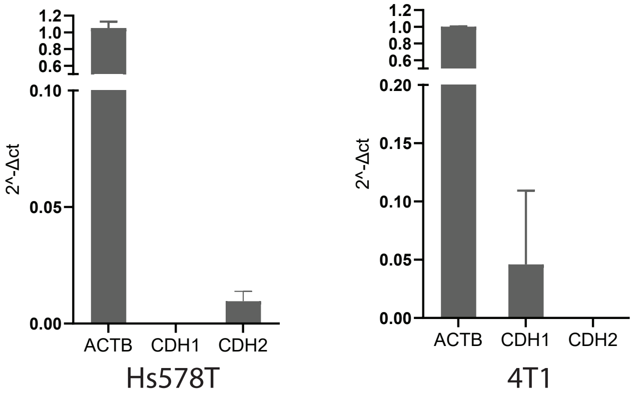

Figure S2. **mRNA expression of E-caherin (CDH1) and N-cadherin (CDH2) in Hs578T (left) and 4T1 (right).** Hs578T cells do not express any E-cadherin but do express N-cadherin. 4T1 cells express E-cadherin but do not express N-cadherin.

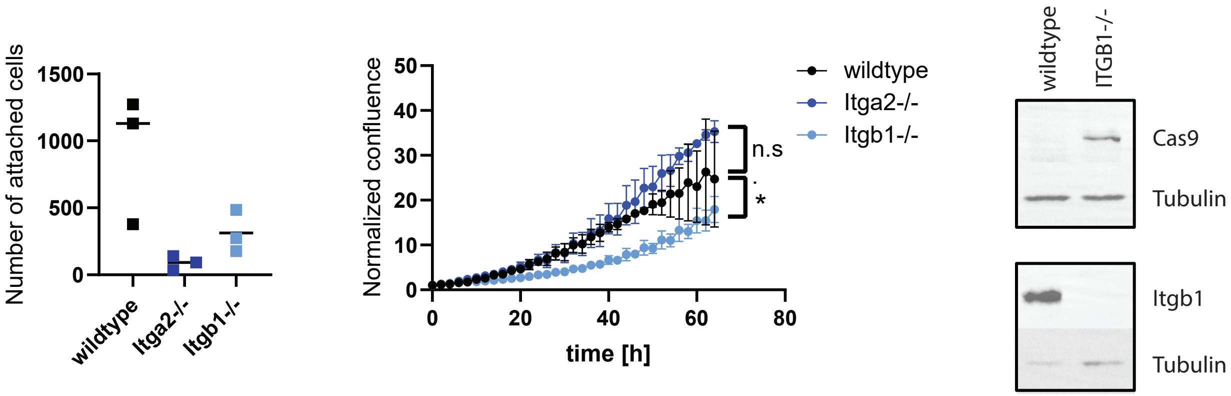

Figure S3. **Deletion of ITGB1 but not ITGA2 affects cell growth of 4T1 cells.** Loss of ITGA2 or ITGB1 leads to reduced collagen binding (left). Loss of ITGA2 or ITGB1 leads to reduced cell growth (middel). Western blot shows complete removal of Itg$\beta$1 from the cells (right).

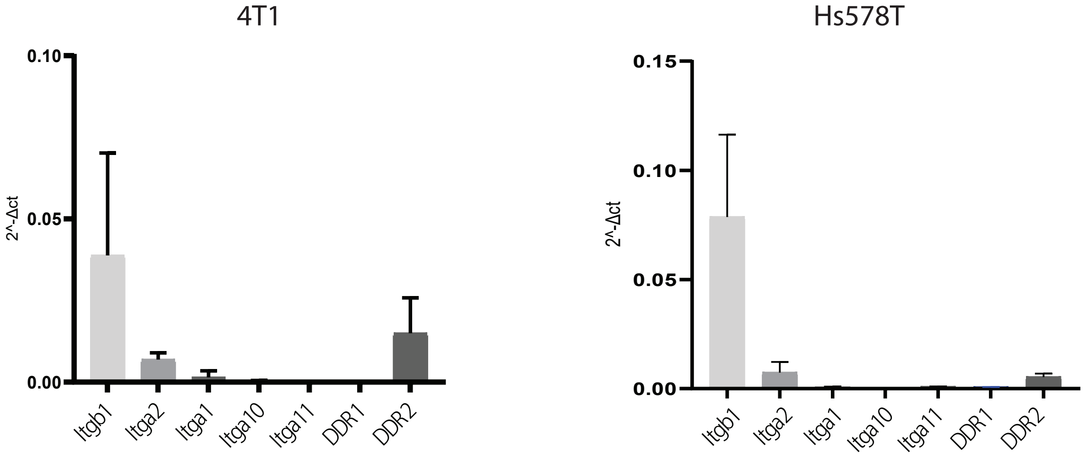

Figure S4. **mRNA expression of collagen receptors in 4T1 (left) and Hs578T (right)**. In both cell lines, integrin $\alpha 2\beta 1$ is the main collagen-binding integrin. Besides integrins both cell lines also express DDR2 as a non-integrin collagen receptor.

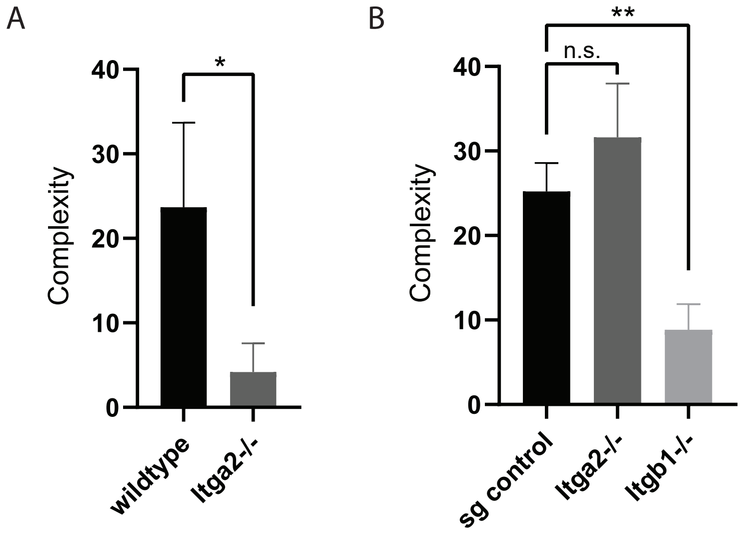

Figure S5. **Complexity diminishes for cells with Integrin knockouts**. **A.** Complexity of 4T1 tumoroids with normal integrin expression (wildtype) and with Itg$\alpha$2 knockout **B.** Complexity of Hs578T tumoroids with normal integrin expression, with Itg$\alpha$2 and with Itg$\beta$1 knockout

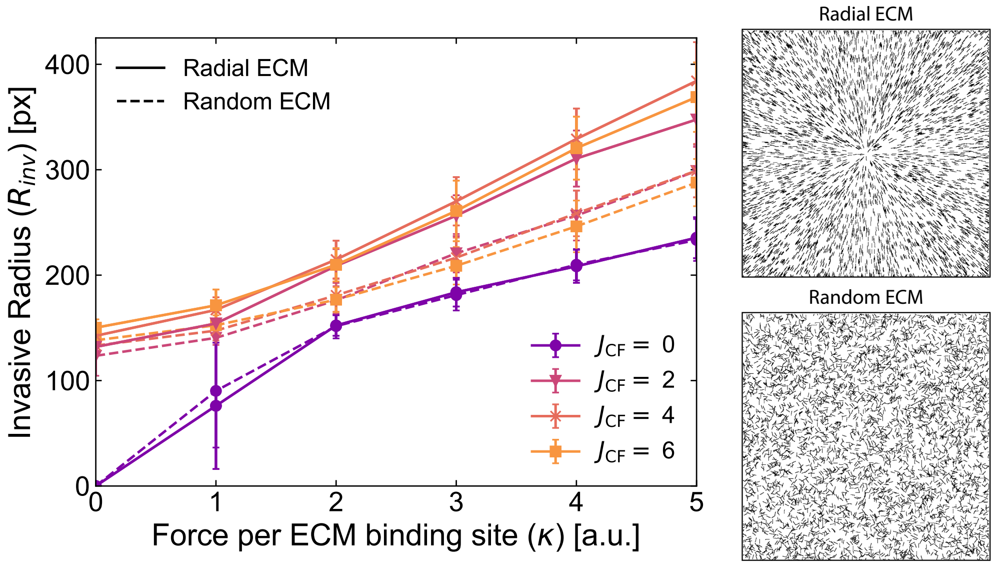

Figure S6. **Invasive radius in two different ECM topologies** The invasive radius is measured for two different ECM topologies, namely a radial network (solid lines) and a random network (dashed lines). The cells in the radial topology migrate further when cell-matrix adhesion ($J_{\mathrm{CF}} > 0$) is present. In absence of any cell-matrix adhesion ($J_{\mathrm{CF}} = 0$), the ECM topology does not play a role.

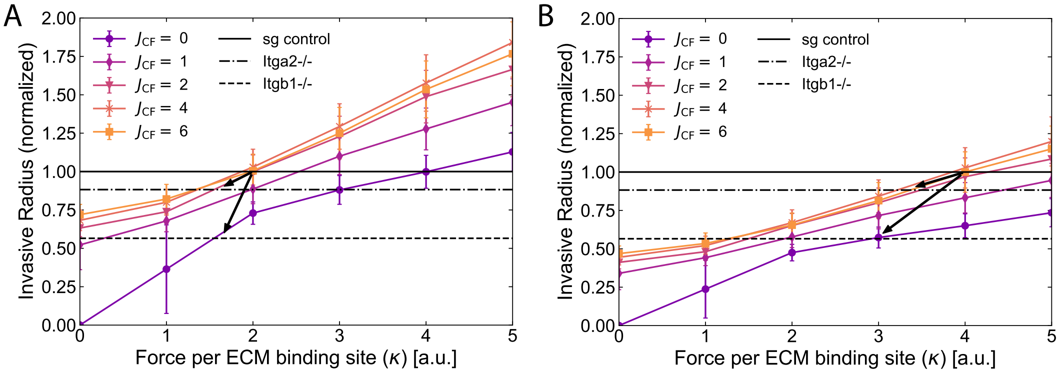

Figure S7. **Comparison between the experiments and the simulations for different reference states. A.** Results for the reference state $\kappa = 2, J_{\mathrm{CF}} = 2$. **B.** Results for the reference state $\kappa = 4, J_{\mathrm{CC}} = 6$. The conclusions do not depend on the chosen reference state. For all reference states, the Itg$\alpha$2$^{-/-}$corresponds to a decrease in the cell-matrix binding while the Itg$\beta$1$^{-/-}$corresponds to a complete KO of cell-matrix binding.

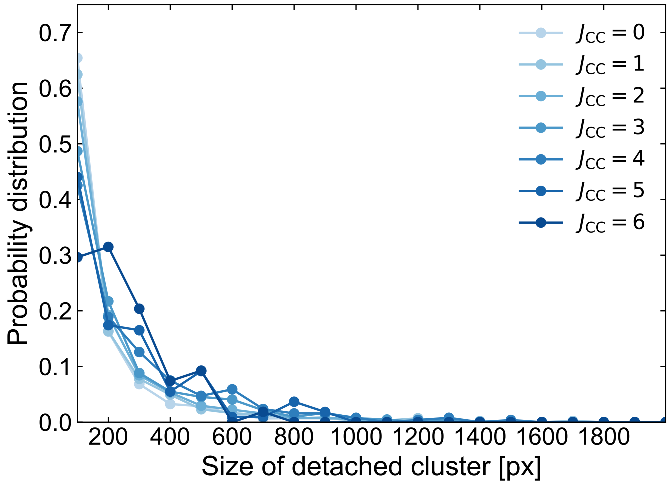

Figure S8. **Multicellular clusters become more dominant as cell-cell adhesion strength increases.** Probability distribution of the detached cells for varying $J_{\mathrm{CC}}$. We kept the cell-matrix adhesion strength fixed at $J_{\mathrm{CF}} = 4$. The average size of a cell is on average 100 pixels.

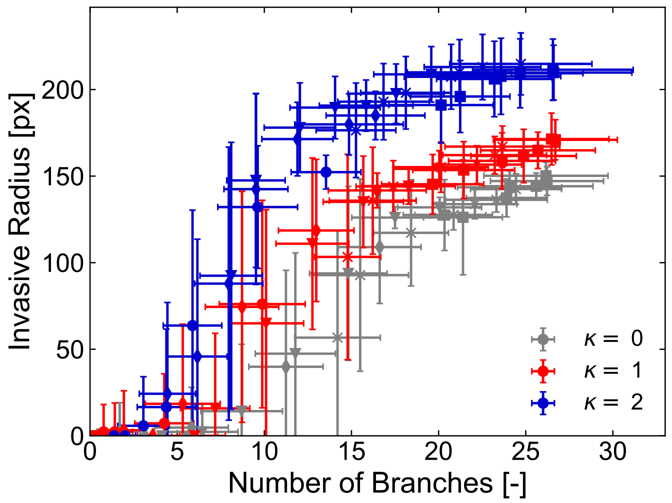

Figure S9. **Collapse of data for varying cell-cell and cell-matrix adhesion holds for different values of $\kappa$.** Invasive radius as a function of the number of branches for the simulations with varying $\kappa$. The data for all values of $J_{\mathrm{CC}}$ and $J_{\mathrm{CF}}$ are colored the same. The observed collapse of the data indicates that knowledge on the specific values of $J_{\mathrm{CC}}$ and $J_{\mathrm{CF}}$ are not necessary, whereas the traction force per ECM binding site ($\kappa$) has a distinct role. Increasing $\kappa$ consistently leads to a larger invasive radius.

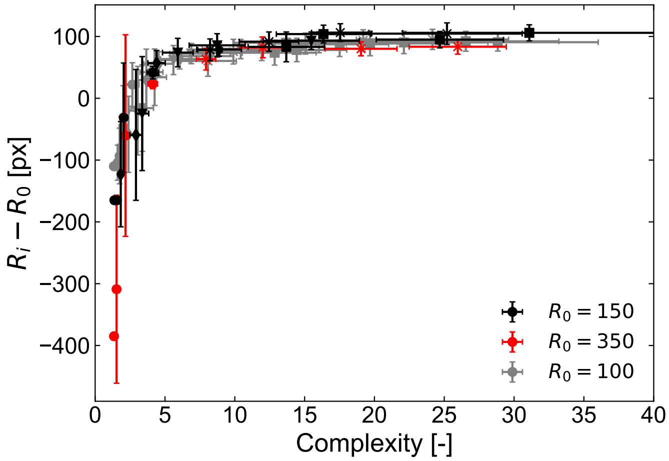

Figure S10. **Data for varying sizes of the primary tumor core collapses** $R_{inv} - R_0$ measures the relative displacement of the most invasive cell with respect to the original tumor radius $R_0$. The data collapses onto a single curve when $R_{inv} - R_0$ is correlated with the complexity of the primary tumor. This suggests that the collapse for data for varying $J_{\rm CC}$ and $J_{\rm CF}$ is independent of the initial size of the primary tumor core. The simulation for $R_0 = 150$ and 350 are for 10 independent runs.

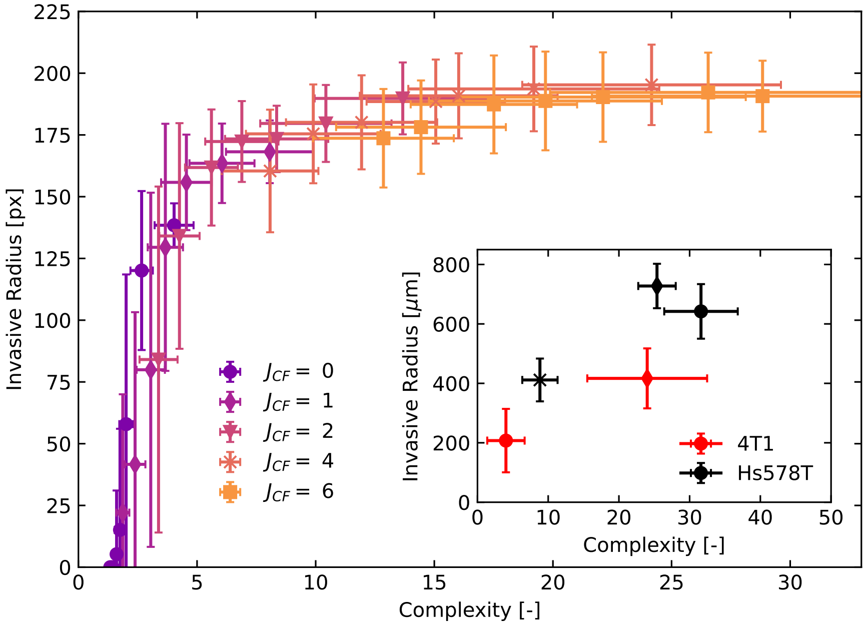

Figure S11. **Direct correlation between the invasive radius and complexity is more difficult to extract from experiments** Invasive radius as a function of the complexity of the primary tumor for varying $J_{\rm CC}$ and $J_{\rm CF}$. The data collapses in the same way as for the number of branches in the simulations, but the same trend is not observed in the experiments.

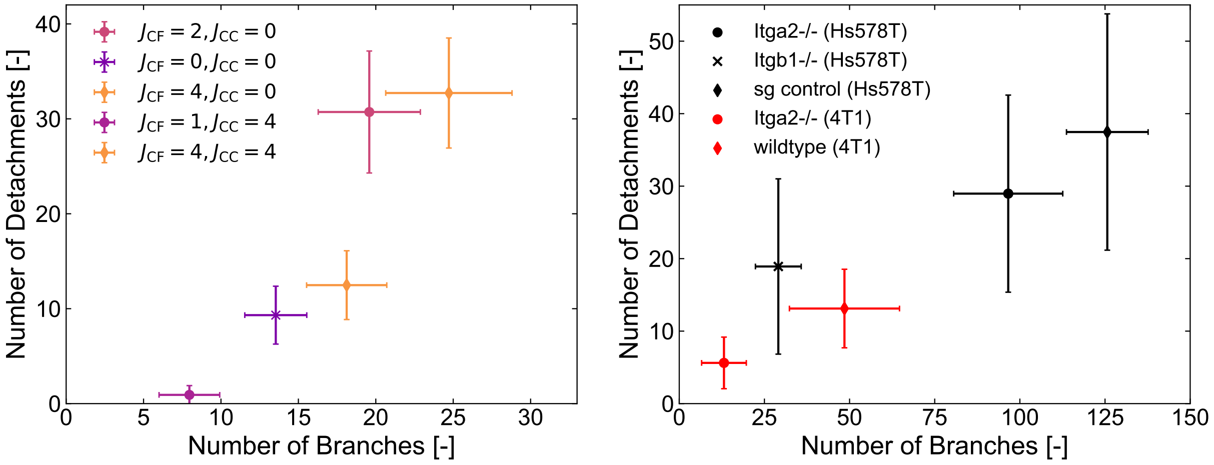

Figure S12. **Correlation between the number of detachments and the number of branches** For the simulation data (left), the results are plotted for the parameter mapping between the experiments and the simulations (found in section A and B). Each line in the legend corresponds to the experimental equivalent in the right panel. The experimental data suggests that the predicted correlation could be present, but the characterization of the number of detachments reliably between the different biological replicates is challenging.
